# Single-cell DNA and RNA sequencing reveals the dynamics of intra-tumor heterogeneity in a colorectal cancer model

**DOI:** 10.1101/616870

**Authors:** Hanako Ono, Yasuhito Arai, Eisaku Furukawa, Daichi Narushima, Tetsuya Matsuura, Hiromi Nakamura, Daisuke Shiokawa, Momoko Nagai, Toshio Imai, Koshi Mimori, Koji Okamoto, Yoshitaka Hippo, Tatsuhiro Shibata, Mamoru Kato

## Abstract

Intra-tumor heterogeneity (ITH) encompasses cellular differences in tumors and is related to clinical outcomes, such as drug resistance. However, little is known about the dynamics of ITH, owing to the lack of time-series analysis at the single-cell level. We performed single-cell exome and transcriptome sequencing of 200 cells and investigated how ITH is generated from one single cell in a mouse colorectal cancer model. The ITH of the transcriptome increased after transplantation from cultured organoids, while that of the exome decreased. Mutations generated in the culture did not greatly change at the transplantation at the bulk-cell level. The RNA ITH increase was due to the emergence of new transcriptional subpopulations. In contrast to the initial cells expressing mesenchymal-marker genes, new subpopulations repressed these genes at transplantation. Analyses of colorectal cancer data from The Cancer Genome Atlas revealed a high proportion of metastatic cases in human subjects with expression patterns similar to the new cell subpopulations in mouse. These results suggest that the birth of transcriptional subpopulations may be a key for adaptation to drastic micro-environmental changes when cancer cells have sufficient genetic alterations at later tumor stages. This study revealed an evolutionary dynamics of single-cell RNA and DNA changes in tumor progression, giving insights into the mesenchymal-epithelial transformation of tumor cells at metastasis in colorectal cancer.

## Background

It is well established that cancer is pathologically composed of different types of cells [1]; however, intra-tumor heterogeneity (ITH) has only been recently addressed at the genomic level [2]. ITH is clinically important. For example, elevated copy-number heterogeneity is related to an increased risk of recurrence or death in non-small-cell lung cancer [3]. High levels of ITH ultimately provide the seeds for the emergence of anti-cancer drug resistance [4]. High levels of genetically-characterized heterogeneity in Barrett’s esophagus are associated with incidence of esophageal adenocarcinoma [5].

ITH essentially stands for the cellular differences in tumor tissue arising from genetic changes, called clonal evolution, or non-genetic changes, such as cancer stem cells and simple transcriptional responses to the environment. In clonal evolution, as in Darwinian evolution, cancer cells with advantageous genetic mutations evolve into different types of cancer cells [6]. In contrast, cancer stem cells, like normal stem cells, produce a variety of differentiated daughter cells that constitute phenotypically distinct cancer cells without genetic differences through epigenetic and the resultant transcriptional mechanisms [7, 8].

A flood of studies have addressed ITH through the variant allele frequencies (VAFs) of tumor cells in bulk, which are calculated from sequence reads with variants identified through next-generation sequencing (reviewed in [2, 9]). In this bulk-cell sequencing approach, the presence of minor clones is often reflected on lower VAFs than the VAF of the major clone [10]. However, this bulk-cell DNA sequencing approach is limited in revealing genetic ITH because it only infers the presence of clones, not directly observing individual cells. In addition, the bulk-cell approach is generally not suitable to resolve transcriptomic ITH, where transcript mixtures from different cells are sequenced.

Single-cell sequencing is a powerful technology for investigating ITH by identifying genomic alterations and distinct transcriptomic states in single tumor cells [11-19]. For example, in clinical samples of glioblastoma, single-cell RNA sequencing showed that individual tumor cells vary in terms of their degree of stemness-related gene expression from extremely stem-like to differentiated states [13]. Additionally, the existence of cancer stem cells that continuously differentiate into astrocyte- and oligodendrocyte-like cells has been demonstrated in oligodendrogliomas by single-cell RNA sequencing [14]. Single-cell DNA sequencing has also been applied to breast cancer samples to evaluate ITH originating in genomic DNA, leading to the suggestion of stepwise/sweepstake or gradual evolution of cancer cells from single nucleotide variation (SNV) data [11, 12, 20]. However, these types of ITH and their respective evolutionary mechanisms are based on snapshot data at one time-point. Furthermore, either RNA or DNA was solely examined. It is necessary to address both RNA and DNA over time for the full elucidation of tumor evolutionary dynamics associated with ITH.

Mouse models are more useful than human clinical samples for examining changes in genomic and transcriptomic states over time. In a breast tumor xenograft mouse model, single-cell DNA sequencing of serially passaged samples identified tumor cell subpopulations and suggested that tumor cells in the same initial state followed the same evolutionary trajectory [21]. In the present study, we employed a modified version of the mouse colorectal cancer model that we previously established [22] and sequenced both single-cell DNA and RNA. We thus investigated how ITH based on the exome and transcriptome changes over time at the single-cell level.

## Results

### Colorectal cancer mouse model

The colorectal cancer mouse model was established by knocking down *APC* expression in normal epithelial cells taken from mouse intestinal crypts using short hairpin RNA (shAPC; **Fig. 1A**) [22]. In the previous system, we used bulk cells from a tissue for culture; however, in this study, we cultured organoids from *one single cell* so that heterogeneity observed in these cultures could not be confused with heterogeneity originating from the knock-down efficiency or intestinal crypts [23].

**Fig. 1.**
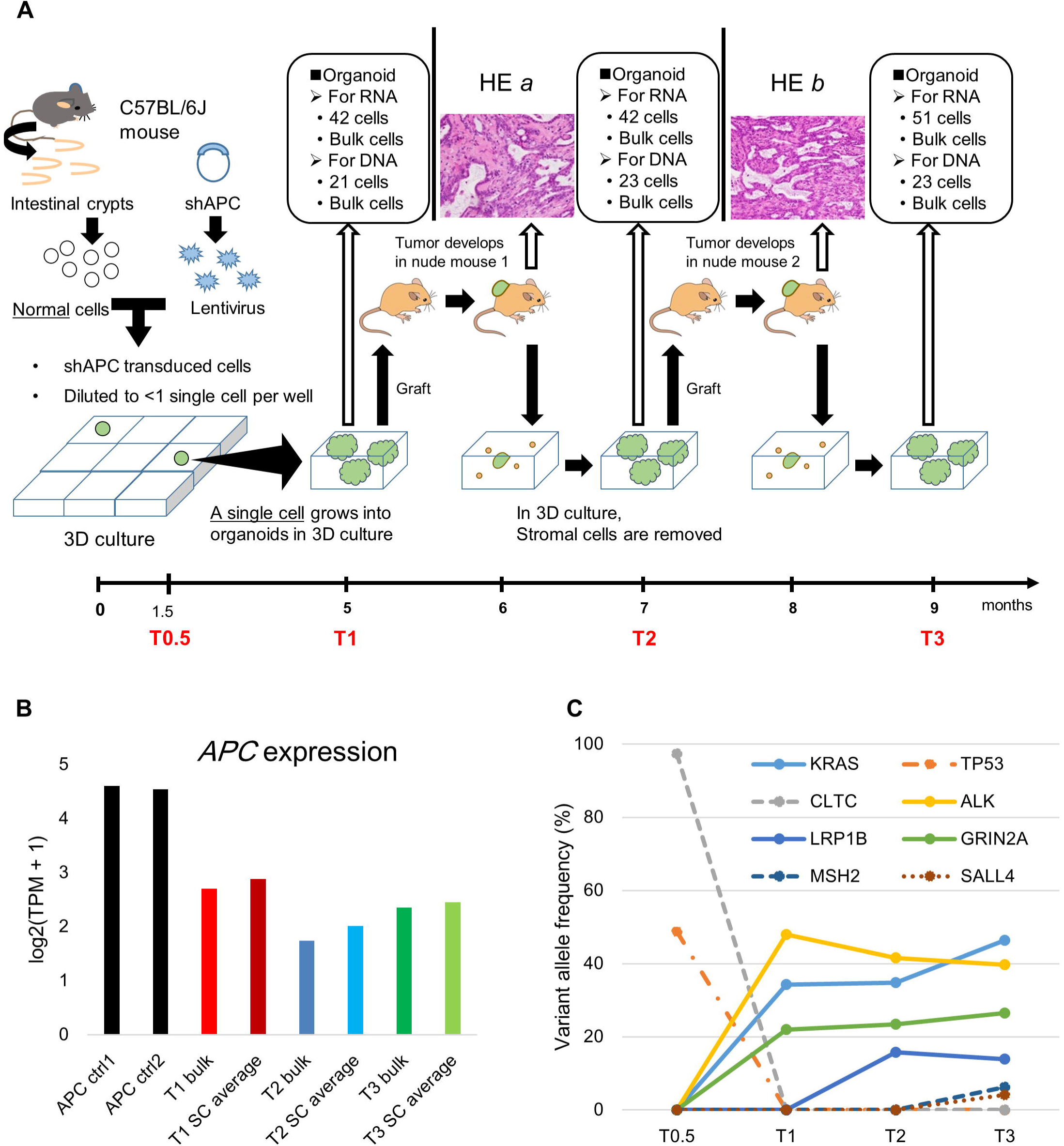
The mouse model. (A) The experimental procedure and HE staining of subcutaneously transplanted tumors. *One single cell* was 3D-cultured in a compartment in a 96-well plate, and single-cell derived organoids were taken to separate single cells. RNA and DNA were separately extracted from the different single cells of multiple organoids and then sequenced. The numbers of cells for RNA and DNA sequencing in boxes are those obtained after quality control of data. (B) The *APC* gene expression from bulk- and single-cell RNA sequencing. “APC_ctrl” indicates control samples that were cultured in our 3D culture system and derived from normal cells without *APC* knockdown at bulk-cell sequencing. “SC average” indicates the average across single cells at each time point. (C) Variant allele frequencies of mutations found in the significantly mutated genes of colorectal cancer by bulk-cell DNA sequencing. See Additional file 1: **Figure S1** for the annotations of the mutations.

We grew organoids for a period of five months so that a single cell having only artificial *APC* intervention could naturally obtain mutations to transform into tumor cells. Cultured cells were subcutaneously transplanted into a nude mouse. One month after transplantation, the mouse was sacrificed, and the tumor was harvested; half of the tumor tissue was re-cultured in our three-dimensional (3D) culture system for one month for the removal of stromal cells. Using half-samples preserved the same genetic lineage over time. The process was repeated once more. Cells were sampled immediately before the first transplantation at time point T1 and at two time points T2 and T3 following the first and second transplantations, respectively (**Fig. 1A**). We sequenced single-cell RNA and DNA separately taken from the different single cells of multiple organoids, which descended from one single cell. We also obtained a DNA sample before T1, at T0.5, 1.5 months after culture initiation (**Fig. 1A**).

Hematoxylin-eosin (HE) staining revealed that subcutaneously transplanted organoids formed tumors consisting of both glandular and non-glandular structures (HE *a* and *b* in **Fig. 1A**). Glandular components in HE *a* were mainly lined with single-layered epithelia, while those in HE *b* were characterized by increased multi-layered regions, loss of cellular polarity, and nuclear enlargement. Non-glandular components had a stromal/medullary structure consisting of spindle-shaped or round to polygonal cells, were characteristically gelatinous/fibrous, and had an abundance of fibrous stroma.

The *APC* expression was decreased in the *APC* knockdown samples (**Fig. 1B**). Out of the 31 significantly mutated genes (excluding *TTN*) defined by The Cancer Genome Atlas (TCGA) colorectal cancer study [24], we found two mutations in *KRAS* and *TP53* by bulk-cell DNA sequencing in our model (**Fig. 1C**), though the *KRAS* mutation was located outside of, but close (9 bps) to, an exon and the position was evolutionary conserved as much as exons (Additional file 1: **Figure S1**). The *KRAS* mutation occupied only a small fraction (2.5%) of the population at T0.5 but increased to 46.4% at T3. Additionally, we found nonsynonymous mutations in six, *CLTC, LRP1B, ALK, GRIN2A, MSH2*, and *SALL4* out of the cancer-related genes in COSMIC Gene Census [25] (**Fig. 1C**). It seems that a major clone with the *TP53* mutation were extinct between T0.5 and T1 and instead minor clones with the *ALK* and *KRAS* mutations became major clones.

### Single-cell transcriptome analysis

We checked various indices of single-cell transcriptome data to filter 42, 42, and 51 cells out of the 50 T1, 43 T2, and 52 T3 cells, respectively (Additional file 1: **Figure S2**). The median (± inter-quartile range) number of mapped reads, mapping rate, and number of expressed genes across selected cells were 6.2 × 10^6^ (± 2.0 × 10^6^), 61.9% (± 5.39%), and 3814 (± 889.5), respectively. There was a strong correlation between gene expression levels in the bulk sequencing data and average expression levels across single cells (Additional file 1: **Figure S2**; *R*^2^ = 0.9). The expression levels of housekeeping genes (GAPDH and mtATP6) [26] were well reproduced across T1, T2, and T3 (Additional file 1: **Figure S2**).

A principal component analysis (PCA) plot of cells based on expression levels revealed increased diversity from T1 to T2 (**Fig. 2A**). This was quantitatively confirmed by the diversity index (distance from the centroid in the PCA space) (**Fig. 2B**). In the plot, T2 and T3 cells partly overlapped but were separate from T1 cells. We identified genes whose expression levels varied greatly across cells at each time point; that is, these genes had high corrected coefficient of variation (c*CV*) values (Additional file 1: **Figure S3**), and were thus referred to as highly variable genes. There were eight, 14, and 16 highly variable genes at T1, T2, and T3, respectively, reflecting an increase in variability from T1 to T2.

**Fig. 2.**
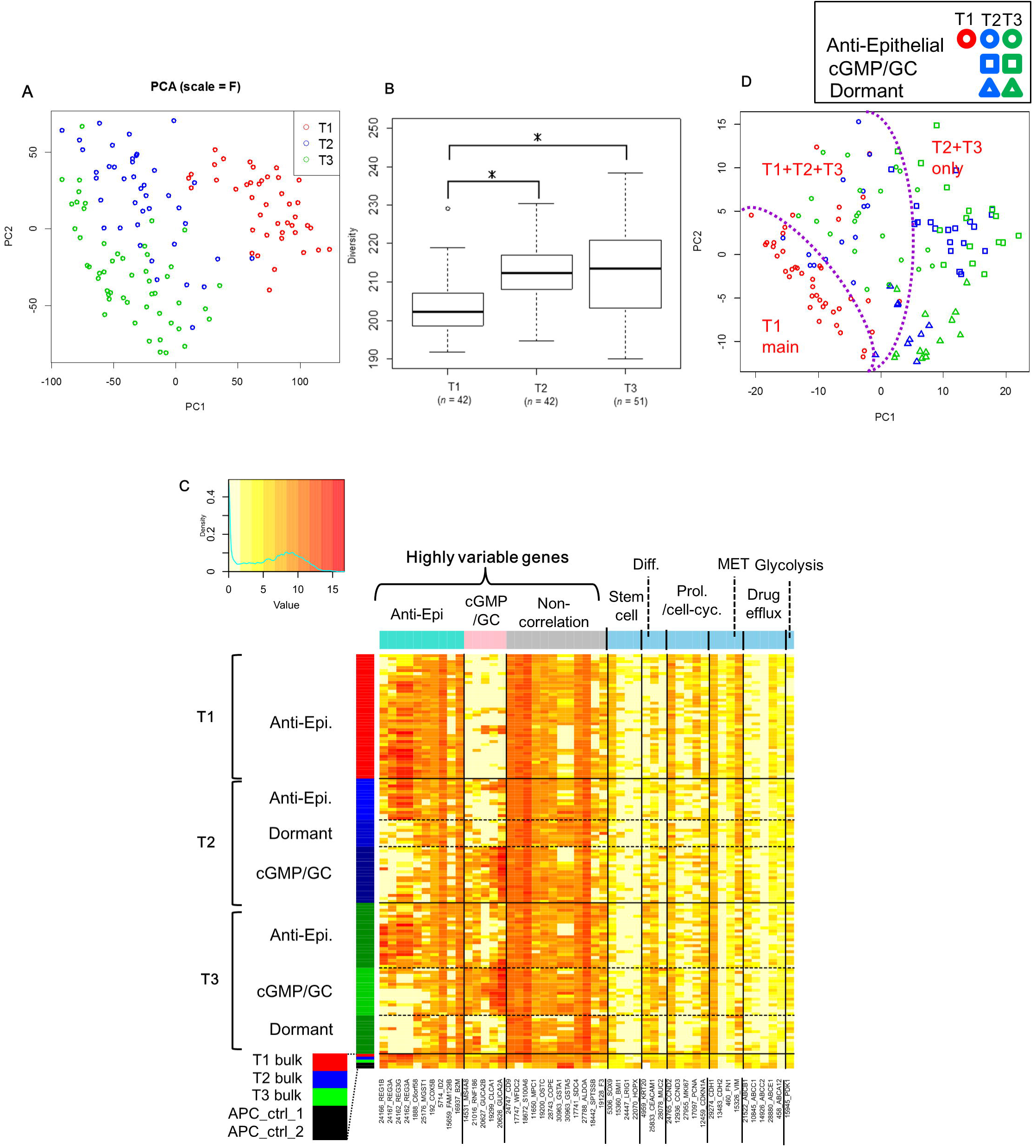
Transcriptome analysis. (A) PCA plot of single cells based on expression levels (genes with TPM ≥ 10 in at least one cell). T1, at the time of 3D culturing; T2 and T3, after the first and second transplantations, respectively. (B) Euclidean distance from the centroid in the PCA space (using full dimensions). \**P* < 0.01 (two-sided Wilcoxon rank sum test). (C) Heatmap of gene expression levels (in TPM). The rows represent single cells or bulk-cell samples (in the bottom), and the columns represent highly variable genes and several types of marker genes. The cell and gene groups were determined as shown in Additional file 1: Figure S4. The red, blue, and green codes in the rows correspond to T1, T2, and T3. “Diff.” and “Prol./cell-cyc.” represents differentiation and proliferation/cell cycle. “APC_ctrl” indicates control samples that were cultured in our 3D culture system and derived from normal cells without *APC* knockdown. (D) PCA plot of cells grouped based on expression levels of highly variable genes.

A cluster analysis of highly variable genes identified three gene groups (Additional file 1: **Figure S4**); expression levels were correlated within two of the groups, but not within the third group. Gene set enrichment analysis showed that one of the correlated groups was associated with negative regulation of keratinocyte differentiation (referred to as Anti-Epithelial genes) (*P* = 3.80 × 10^−3^), whereas the other was associated with positive regulation of cGMP and guanylate cyclase (GC) activity (referred to as cGMP/GC genes) (*P* = 1.30 × 10^−3^), which are known to be associated with negative regulation of β-catenin signaling and matrix metalloproteinase activity in colorectal cancer [27, 28].

A heatmap generated from the cluster analysis revealed that T1 cells were relatively homogenous and formed one group that highly expressed Anti-Epithelial genes but showed negligible expression of cGMP/GC genes (**Fig. 2C**). This group was therefore termed Anti-Epithelial. In addition to an Anti-Epithelial cell group, two new groups appeared at T2: one showing the opposite pattern, repression of Anti-Epithelial and activation of cGMP/GC gene expression, referred to as the cGMP/GC cell group; the other showed repression of both Anti-Epithelial and cGMP/GC genes, referred to as the Dormant group for the marker analysis described below. Notably, as shown in the heat map, bulk-cell sequencing analysis alone could not have identified these cell groups, where their distinct expression patterns were offset by bulk-cell expression levels (labeled as T1, T2, and T3 bulk in **Fig. 2C**).

Using a PCA plot based on highly variable gene expression, we confirmed that T1 cells were relatively homogeneous and T2 cells showed similar grouping to T3 cells (**Fig. 2D**). Looking at the PCA closer, we found that cells of the Anti-Epithelial group seemed close together across all time points, but seemed to form two groups—i.e., T1 main (referred to as T1 main) and T1/T2/T3 mixture (T1+T2+T3). The cGMP/GC and Dormant groups seemed close together and contained T2 and T3 only (T2+T3 only).

### Marker gene expression

We examined the expression of several types of marker genes. We first looked at proliferation/cell cycle markers (Additional file 1: **Figure S5**) and performed PCA to summarize the multiple expression levels (**Fig. 3A**). Remarkably, most cells in the Anti-Epithelial group at T1 expressed high levels of proliferation- and cell cycle-related genes according to the PCA loading plot. In contrast, nearly all cells in the Dormant group at T3 showed a downregulation of the marker genes (so the cell group was termed Dormant). At T2, about half of the cells showed a downregulation of the proliferation/cell-cycle genes.

**Fig. 3.**
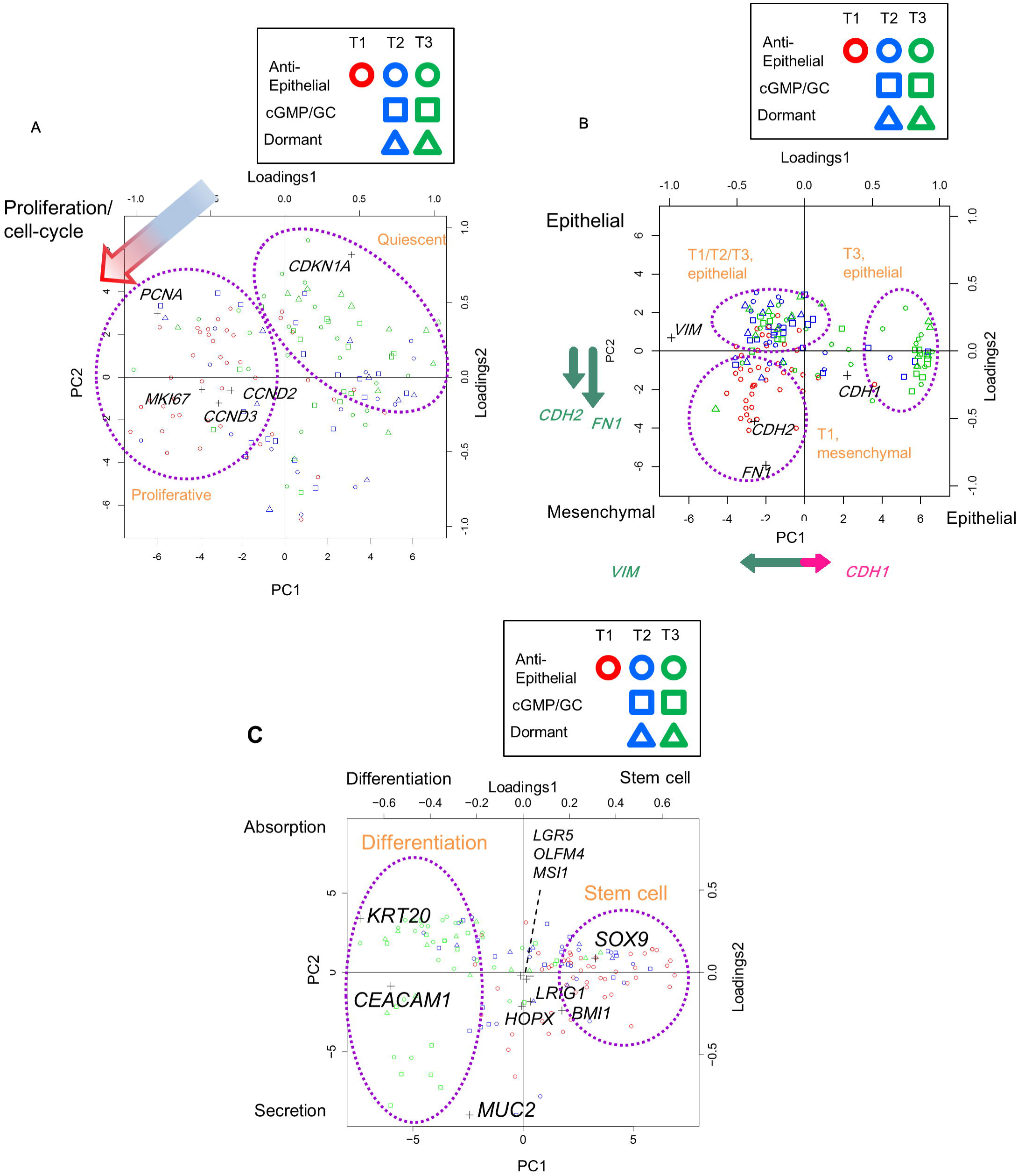
PCA and overlaid loading plots based on expression levels of markers (A) about the proliferation/cell cycle. The arrow indicates the direction from negative to positive markers in the loading plot; cells positioned in that direction in the PCA plot had higher expression levels of positive marker genes. (B) About the epithelial and mesenchymal. The arrows along the *x* and *y* axes represent projected loadings in the loading analysis, where cells positioned in that direction in the PCA plot had higher marker gene expression levels. (C) About stem cell and differentiation.

We next examined epithelial and mesenchymal markers (Additional file 1: **Figure S5**). A PCA plot of the markers showed that expression of mesenchymal cell-related genes decreased with time (T2 and T3), with cells forming two groups (**Fig. 3B**): one (upper left) overlapping with some T1 Anti-Epithelial cells with decreased mesenchymal N-cadherin (*CDH2*) and fibronectin (*FN1*) levels; the other (middle right) group was composed only of T3 cells with decreased mesenchymal vimentin (*VIM*) and increased epithelial E-cadherin (*CDH1*) levels. These results suggest a similarity between the processes occurring from T1 to T2 and mesenchymal-epithelial transition (MET).

Stem cell and differentiation markers showed that over time, cells generally expressed more differentiation than stem cell markers (**Fig. 3C**; Additional file 1: **Figure S5**), though a remarkable variation across individual cells was also observed. Fractions of cells with the expressions of stem cell markers decreased with time. Among the markers for crypt base stem cells, *SOX9* appeared to be the most influential; *LGR5, OLFM4*, and *MSI1* were not substantially expressed. It seems that with time, cells differentiated into those expressing a marker for absorption cells (*KRT20*) and those for secretion cells (*MUC2*) in the digestive tract.

There was no remarkable change in the expression of drug efflux genes [29, 30] at any time point (**Fig. 2C**), although *ABCB1* expression was slightly lower in the T3 Dormant group (Additional file 1: **Figure S5**) and *ABCE1* was downregulated at T2 and T3. There was variable expression of glycolysis-related gene *PDK1* [30] across all cells, irrespective of groups (**Fig. 2C**; Additional file 1: **Figure S5**).

### Single-cell exome analysis

Based on several indices from single-cell exome sequencing (Additional file 1: **Figure S6**), we selected 21, 23, and 23 cells out of the 23 T1, 24 T2, and 24 T3 cells for analysis. On average (expressed as the median [± inter quartile range] across selected cells), the number of mapped reads was 1.2 × 10^8^ (± 2.2 × 10^7^), mapping rate was 76.6% (± 4.9%), coverage with > 0 depth regions was 76.9% (± 34.2%), average depth was 43 (± 34.5), Gini coefficient was 0.85 (± 0.15), allelic drop-out (ADO) rate was 47.0 (± 36.1), and number of called SNVs was 462 (± 313.5). The false positive rate in single-cell sequencing was estimated to be 0.1–1.1 × 10^−7^ per chromosomal site, based on normal intestinal tract tissue samples from two mice and four single cells obtained from one of these samples (**Additional file 1: Supplementary Results**). We compared the fractions of single cells with SNVs to the variant allele frequencies (VAFs) of the bulk-cell sequencing; in theory, the single-cell fractions should be equal to half of the VAFs. We confirmed a good concordance between these variables, although the cell fractions were slightly lower than those expected from bulk VAFs (Additional file 1: **Figure S6**).

We first examined the bulk-cell sequence data. The T0.5 tissue had much fewer SNVs than the later stages (**Fig. 4A**), which suggests that DNA heterogeneity only weakly appeared soon (1.5 months) after culture initiation. The numbers of SNVs increased markedly from T0.5 to T1, a five-month period (**Fig. 4A**). Although these numbers decreased slightly at T2 before recovering at T3, they were all mostly saturated at T1, T2, and T3. Thus, new SNVs were largely generated from T0.5 to T1, and most of these SNVs remained in the genome after T1 at the bulk-cell level (**Fig. 4B**).

**Fig. 4.**
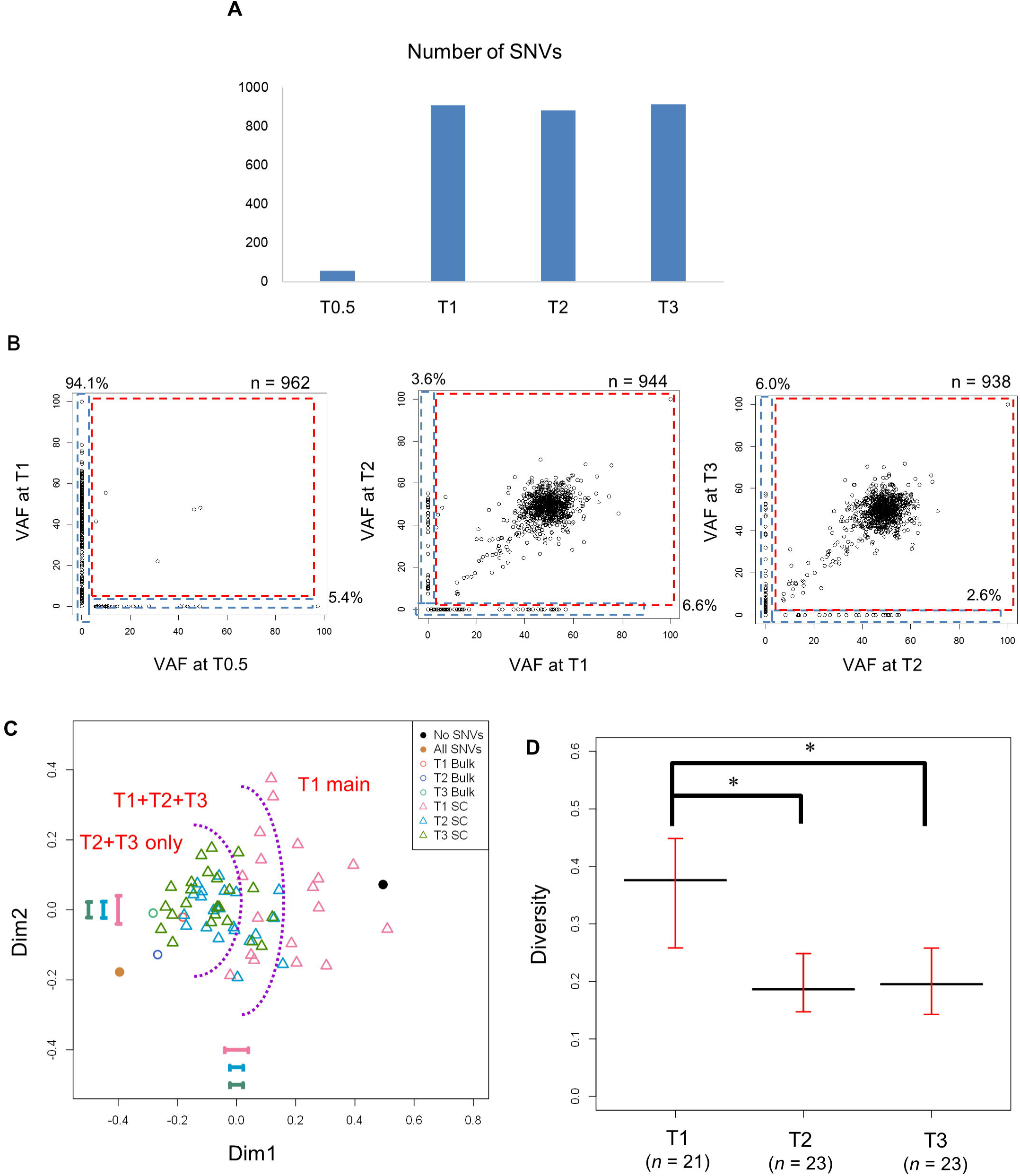
Exome analysis. (A) Number of SNVs called in bulk-cell sequencing. (B) Comparison of VAFs of SNVs called in bulk-cell sequencing at successive time points. One point indicates one SNV. Numbers represent the number of points. (C) MDS plot based on single-cell exome sequencing. “No SNV” and “All SNV” represent sequences with no SNVs and with SNVs at all sites, respectively, which were artificially generated as a reference. Error bars represent the standard deviation for each dimension calculated with a bootstrapping approach that took into account ADO rates. (D) Median Euclidean distance from the centroid over cells in the MDS space. The black and red bars represent the observed value and 95% confidence interval calculated with the bootstrapping approach. \**P* < 0.05 (bootstrapping test).

We then used single-cell sequencing data to draw a multi-dimensional scaling (MDS) plot based on single-cell SNVs at polymorphic SNV sites (defined as SNVs with 10–35% bulk VAFs) (**Fig. 4C**). T1 cells showed the greatest genetic diversity, whereas T2 and T3 cells showed less diversity. This decrease in diversity was confirmed by a statistical significance of the diversity index (average distance from the centroid), where the bias due to ADO rates was taken into account by a bootstrapping test (**Fig. 4D**). Interestingly, this diversity tendency was the complete opposite of the transcriptomic pattern (**Fig. 2A, B**). Although transitional, cells can be classified into three groups composed of T1 cells only (T1 main); T1, T2, and T3 cells (T1+T2+T3); and T2 and T3 cells (T2+T3 only) (**Fig. 4C**), which could correspond to the three groups in the single-cell transcriptome analysis (**Fig. 2D**)

### Association with human cancer

We used 244 TCGA samples of colorectal cancer [24] to identify human samples with gene expression patterns similar to the groups of mouse single cells by MDS (**Fig. 5A**). In the MDS plot, the human samples were separated from the mouse cell groups, but we found that 94 (38.5%), 42 (17.2%), and 13 (5.3%) human samples were respectively close to the Anti-Epithelial, cGMP/GC, and Dormant mouse cell groups. TCGA Anti-Epithelial samples showed enhanced *REG* and repressed cGMP/GC gene expression; TCGA cGMP/GC samples showed the opposite pattern; and TCGA Dormant samples had both repressed *REG* and GC-related gene expression (**Fig. 5B**). TCGA cGMP/GC and TCGA Dormant samples tended to be more closely associated with metastasis than those with patterns similar to the Anti-Epithelial group (two-sided Fisher’s exact test *P* = 0.04; **Fig. 5C**). We determined that our mouse tumor cells were molecularly similar to cells of human colon adenocarcinoma classified as high microsatellite instability type (Additional file 1: **Figure S7**; Additional file 1; **Figure S8**; Additional file 1: **Supplementary Results**).

**Fig. 5.**
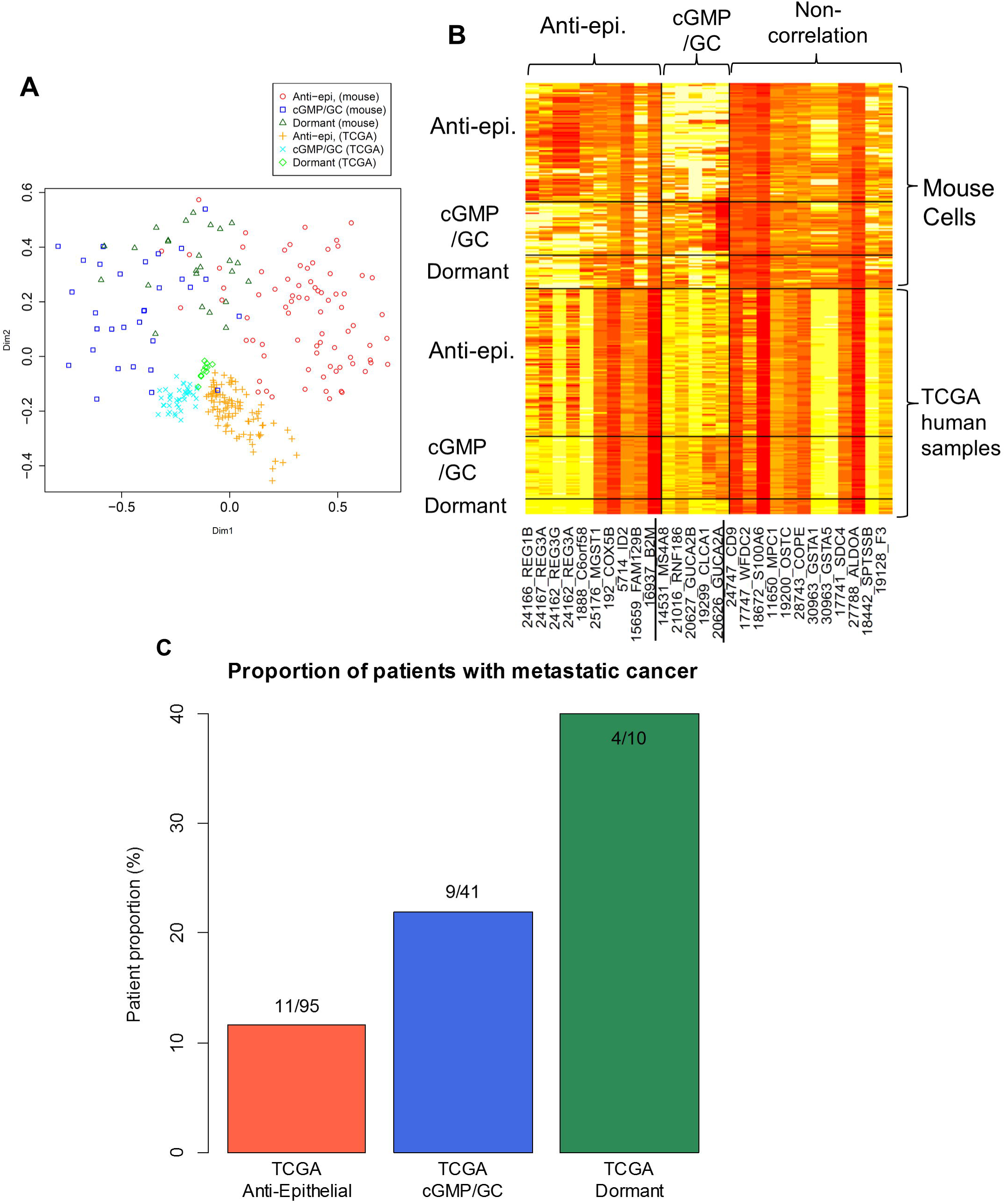
Analysis of TCGA human samples with gene expression patterns similar to mouse cell groups. (A) MDS plot of mouse single-cell samples and such TCGA samples on the basis of a similarity of gene expression patterns. (B) Heatmap of the samples. Genes are highly variable genes shown in **Fig. 2C**. (C) The fraction of patients with metastatic tumor in TCGA samples with expression patterns similar to mouse cell groups

## Discussion

Our mouse model was close to human colon adenocarcinoma of the MSI-high hyper-mutation type. In this model, once cancer cells accumulate a sufficient number of genetic alterations (SNVs/indels), they may be able to adapt to drastic environmental changes, such as the shift from a 3D culture to a live mouse, by only altering their transcriptional profiles without further genetic changes. Such transcriptional adaptation may cause the generation of new subpopulations, leading to increased transcriptional heterogeneity. Meanwhile, genetic heterogeneity decreased, possibly as a result of microscale natural selection that occurred during the environmental transition. Though expected, it is nonetheless surprising to see that this diversity was indeed generated from *one single cell*.

T1 cells had the saturated number of genetic mutations, expressing active cell-cycle, mesenchymal, and stem cell markers. Thus, the cells are considered as those at a late tumor stage when they move out from the niches or microenvironment of intestinal crypts [31]. Moreover, the emergence of the Dormant and cGMP/GC groups at T2 and T3 was associated with metastasis in the analysis using TCGA human samples. Therefore, our observation that cells lose their mesenchymal-like phenotype and acquire epithelial-like characteristics after subcutaneous transplantation may be analogized to MET during metastasis, though this implication should be tested by further investigations of clinical samples including single-cell sequencing of TCGA samples.

Classically, cells that generate a tumor by subcutaneous transplantation are called tumor-initiating cells or cancer stem cells (CSCs) [30]. In this classical model, it is expected that differentiated cells die while CSCs can survive at the start of subcutaneous transplantation and 3D culture; then, CSCs regenerate differentiated cells. We initially expected that fractions of cells expressing stem cell markers increased over the serial transplantation, because we simply thought that it is CSCs, not differentiated cells, that can survive at transplantation. However, our observation was the opposite. CSCs that efficiently generate differentiated cells may be more adaptive for the merit of obtaining mutual benefits between different types of cells. Alternatively, contrary to the classical expectations, CSCs may not necessarily express high levels of stem cell markers: the Dormant and cGMP/GC cells with low expression levels of stem cell markers may be also CSCs or tumor-initiating cells that survived at the transplantation and 3D culturing.

Recently, more fine-scale single-cell sequencing technology, such as 10X/Drop-Seq, has emerged for RNA-seq, enabling researchers to capture tens of thousands of cells. Although the number of cells we addressed was relatively small compared to that technology, we believe that we successfully captured a major part of the heterogeneity constructed by cell clones, constituting as small as ∼2% (an inverse number of 42, 42, and 51 cells at T1, T2, and T3) of the tumor cell population. Nevertheless, 10X/Drop-Seq will be needed to investigate rarer cells.

We demonstrated that time-series ITH analysis by single-cell DNA and RNA sequencing for a mouse model is able to deepen our understanding of the evolutional processes of cancer cells and raise issues on CSCs from the genomic and transcriptomic viewpoints. The birth of transcriptional subpopulations of cells may be a key for adaptation to drastic micro-environmental changes when cancer cells have sufficient genetic alterations at later tumor stages. It will be crucial to examine how such genetic changes accumulate in the earlier stages of tumorigenesis and how transcriptional subpopulations develop to increase malignancy in the further later stages of tumor progression.

## Materials and Methods

### Ethics approval and consent to practice

Animal studies were carried out according to the Guideline for Animal Experiments established by the Committee for Ethics in Animal Experimentation of the National Cancer Center (T10-033-M05), which meets the ethical standards required by law and guidelines for animal experimentation in Japan. All sacrificed mice were anesthetized by inhalation of isoflurane. And cervical dislocation was used as a euthanasia method.

### Organoid culture of small intestinal cells and lentiviral transduction

C57BL/6J mice and BALB/cAnu/nu immune-deficient nude mice were purchased from CLEA Japan (Tokyo, Japan). The small intestine was harvested from wild-type male C57BL/6J mice at 3–5 weeks of age. Crypts were purified and dissociated into single cells, which were then put into serum-free Matrigel-based organoid culture as previously described [22, 32]. Transduced organoids were maintained in culture medium lacking R-spondin 1. Single cell-derived shAPC-transduced organoids were obtained by limiting dilution of dissociated organoids in a 96-well plate. Organoids composed of 5 × 10^5^ cells were mixed with 200 µl of Matrigel and injected into one side of the dorsal skin of nude mice, while uninjected cells were maintained in 3D cultures for later use.

### Analysis of subcutaneous tumors in nude mice

Palpable tumors from the injection sites were harvested for histological examination or cell culture. Half of the subcutaneous tumors were formalin-fixed, paraffin-embedded, and sectioned at a thickness of 5 µm, followed by HE staining to assess histological features. The other half of the tumors were minced and digested to recover cells as described previously (22), then seeded onto solidified Matrigel to resume organoid culture.

### Single-cell transcriptome and exome sequencing

Cultured mouse organoids derived from a single cell were harvested and treated with Accumax (Innovative Cell Technologies, AM105) to generate a single-cell suspension. To capture cells and extract RNA or DNA from a single cell, the cell suspensions (4.4 × 10^5^ cells/ml) were loaded on a C1 Single Cell Auto Prep System (Fluidigm, C1) with medium-sized (10–17 μm) microfluidic chips for 96 cells. Approximately 1300 cells were applied to each chip. Captured cells were imaged on a BZ-9000 digital microscope (Keyence, BZ-9000) and a visual inspection was performed to determine whether a single cell was captured in each well of the chip. Capture efficiency for a single cell was determined as 71–82%.

For single-cell whole transcriptome (RNA) sequencing, captured cells were lysed on the chip using a C1 Single-Cell Auto Prep Reagent Kit for mRNA Seq (Fluidigm, 100-6201). Full-length cDNA fragments were transcribed and amplified from poly-A RNA in each single cell using the SMARTer Ultra Low RNA kit (Takara Bio, 634832). Tagmentation of cDNA was performed and sequencing libraries were prepared using the Nextera XT DNA sample preparation kit (Illumina, FC-131-1096) according to the manufacturer’s protocol. Up to 52 independent single-cell RNA-seq libraries were prepared for sequencing.

For single-cell DNA sequencing, genomic DNA was prepared from single cells using the C1 Single-cell Auto Prep Reagent Kit for DNA Seq (Fluidigm, 100-7357) and whole genome amplification was achieved by multiple displacement amplification with Phi29 DNA polymerase and the Illustra GenomiPhi v.2 kit (GE Healthcare, 25660032). Amplified genomic DNA (70 ng) was used to generate exome sequence libraries using the Hyper Prep kit (Kapa Biosystems, KK8504), SureSelect Target Enrichment kit (Agilent Technologies, 931171), and SureSelect XT Mouse All Exon v.1 probe (Agilent Technologies, 5190-4642).

### Bulk-cell transcriptome and exome sequencing

Among the cells that were not used for single-cell capture with the C1 system, suspensions of about 200 cells were subjected to whole transcriptome (RNA) sequencing for bulk-cell RNA-seq (T1, T2, and T3 samples). The sequencing libraries were prepared using the same reagents as the single cell RNA-seq. As control bulk cells, normal intestinal crypt epithelial cells from two wild-type mice of the same strain were grown in the 3D culture system for seven days, then harvested and lysed for total RNA preparation using the miRNAeasy Mini kit (Qiagen, 217004). RNA-seq libraries for control bulk RNA were generated using the SureSelect Strand-specific kit (Agilent Technologies, G9691B). Bulk DNA from over 1 × 10^5^ cells was obtained from the cell culture (T0.5 sample, 1.5 months after culture initiation) and the remaining cells in single-cell capture (T1, T2, and T3 samples) using the QIAamp DNA Mini kit (Qiagen, 51304), and 500 ng of DNA were used to construct exome sequencing libraries with the same reagents as the single cell DNA-seq.

### Sequencing

All the sequencing libraries were subjected to paired-end sequencing of 101-bp fragments on the HiSeq2500 DNA sequencer (Illumina, SY–401–2501).

### Transcripts per kilobase million (TPM) calculation for single and bulk cells

The procedure for calculating TPM is summarized in Additional file 1: **Figure S9.** Specifically, sequence reads were quality-filtered and trimmed using PrinSeq [33], and then used as input for quality-check by FastQC (https://www.bioinformatics.babraham.ac.uk/projects/fastqc/). We used the following parameters: --min_len 30 (removing reads ≤30 bases); --min_qual_mean 20 (average read quality ≤ 20); --trim_tail_right 5, --trim_tail_left 5 (trim bases if the 3’ and 5’ end poly A/Ts are ≥ five bases); and --trim_qual_right 20, --trim_qual_left 20 (trim 3’ or 5’ end for read quality ≤ 20). Paired-end reads were mapped to the University of California Santa Cruz mouse genome sequence (mm10) using Bowtie2 [34] built in RSEM [35]. Expression levels (in TPM) were estimated by RSEM using the command rsem-calculate-expression with the parameters --estimate-rspd, --paired-end, --bowtie2, -p 30, and --ci-memory 10192. We removed eight T1 cell samples due to an excessive number of genes (≥ 5,200) with TPM ≥ 10 or with too few unique mapping reads (< 2.2 × 10^6^). We also removed two samples with unique mapping rates that were too low (< 20%) and discarded genes with low expression levels (≤ 10 TPM) across all cell samples, leaving 14,258 out of 32,732 genes for analysis.

### Detection of highly variable genes

To detect genes with variable expression levels across cells, we defined highly variable genes according to the *CV*, corrected in the locally weighted scatterplot smoothing (LOWESS) method using the “lowess” function in R. To fit a single LOWESS curve across all ranges, we divided average expression level data into three ranges: < 4, 4–8.5, and > 8.5. c*CV* values were yielded by dividing *CV* values by the value of the upper variability band (± 1.96 times the standard deviation) of smoothed curve estimated using “loess.sd” in the “msir” package. Because of the large bias in original *CV* values against low average expression levels, only those with c*CV* values > 1.3 and high average expression levels (log_2_ [TPM+1] ≥ 4) were defined as highly variable genes.

### PCA of RNA data

PCA was carried out for gene expression levels (log_2_ [TPM + 1]) without scaling. For the loading analysis of marker genes, we used the following genes; *MKI67* and *PCNA* for positive markers and *CDKN1A* for a negative marker for cell proliferation in colorectal cancer [36]. *CCND2* and *CCND3* for positive markers for cell cycle in this cancer [37]. E-cadherin (*CDH1*) for an epithelial marker; N-cadherin (*CDH2*), vimentin (*VIM*), and fibronectin (*FN1*) for mesenchymal markers [38]. *LGR5, ASCL2, OLFM4, MSI1*, and *SOX9* for crypt base stem cell markers, *HOPX, BMI1*, and *LRIG1* for +4 (position from the crypt base) stem cell markers, *AQP8, CAR1, CEACAM1, KRT20*, and *SLC26A3* for differentiation makers for absorption cells, and *MUC2, SPINK1* for differentiation markers for secretion cells [39].

### Hierarchal clustering, correlation plot, and heatmap analysis

For hierarchal clustering, we used the “hclust” function in the “stats” package of R software, where we calculated the Euclidean distance of expression levels (log_2_ [TPM +1]) of all highly variable genes between cells and used the Ward method for agglomeration. We generated correlation plots of highly variable genes using the “corrplot” function in the R “corrplot” package, where we used the Ward method for agglomeration. We divided genes into three clusters based on these hierarchical clustering results using the “addrect = 3” option. A heatmap was generated using the “heatmap.2” function in the “ggplot2” package. In the heatmap, cells were arranged according to their order in the dendrogram described above and genes were arranged according to their order in the correlation plot of highly variable genes.

### Gene set enrichment analysis

DAVID [40] was used to identify gene ontologies (biological processes) in which genes of an identified group were enriched (*P* < 0.01).

### SNV detection for single and bulk cells

For bulk-cell data, we used a previously described method for SNV/indel calling [41] by cisCall with cell-line/frozen parameters [42], mapping reads to the mouse genome (mm9) by BWA [43]. We filtered out PCR-duplicated reads as well as reads and bases with low mapping and base qualities. The remaining variants were further filtered statistically using Fisher’s exact test to compare fore- and background samples, which came from two different individuals of the same pure C57BL/6J strain. We verified the negligible effects of using a different individual for the background sample (**Supplementary Results**). A series of filters was used to remove suspicious variant calls, such as those related to misalignments. Variants for which allele frequencies were significantly greater than 1% in the binomial test were retained. The procedure is summarized in Additional file 1: **Figure S9**.

For single-cell sequencing data, we called SNVs only at SNV sites called in bulk-cell sequencing data. Specifically, we counted nucleotide bases with high qualities (mapQ ≥ 20, BaseQ ≥ 10) in single-cell sequencing data as well as in background data (same as those used in bulk-cell SNV calling) with the Samtools mpileup function [43, 44]. We then selected variants with the largest *χ*^*2*^ test statistic (with Yates’s correction) among all possible variants at each position to identify those that were most likely to differ between single-cell and background data. We required a variant count ≥ 2 and VAF ≥ 2% for such variants in single-cell data. We then selected variants that overlapped with SNV sites called in bulk-cell data.

### Detecting mutation in cancer-related genes

We investigated nonsynonymous mutations in cancer-related genes contained in Tier1 in COSMIC Cancer Gene Census [25].

### MDS of DNA data and the diversity index

We performed MDS from the cell × site matrix composed of zero and one, which respectively represent the absence and presence of SNVs (both synonymous and non-synonymous SNVs) and NA, which represents an undetermined call due to shallow depth. We assigned zero to non-called sites as the true negative when those sites had depths ≥ 30 and assigned NA to non-called sites when the depth was < 30. We only used SNV sites where a variant was called in at least one cell and the VAFs at the same sites in bulk data ranged from 10–35% (polymorphic) for at least one time point. We removed cells and sites (two each) with too few or too many NAs, yielding 104 sites and 69 cells. Using this 0/1/NA matrix, we calculated the *p*-distance (proportion of different sites) used in molecular evolution without using NA, and then performed MDS.

The diversity index was calculated as the average Euclidian distance from the centroid over cells in the MDS space, where we used up to the sixth dimension because of a sudden drop in the eigenvalues over this dimension. To calculate the statistical significance of differences between cell groups, we used a bootstrapping approach in which we randomly re-sampled cells’ sequences from the 0/1/NA matrix of each cell group 10,000 times and performed the same MDS as in the observed data for each replicate. We then calculated the diversity index for each replicate to determine the 95% confidence interval and standard deviation for each cell group.

### Lorenz curve and Gini coefficients

A Lorenz curve was generated with read depth at each site using the “Lc” function in the “ineq” package of R software. To quantify uniformity, the Gini coefficient was calculated using the “Gini” function in the “ineq” package.

### ADO rate

The ADO rate was defined as the rate of homozygous sites in single-cell samples where the bulk sample was heterozygous (defined as sites where VAFs were 45–55%) at the same nucleotide site. We removed outlier cells with high ADO rates at each time point (one cell each with an ADO rate > 80% at T2 and T3).

### Average copy number

The average copy number (ACN) was calculated as follows:

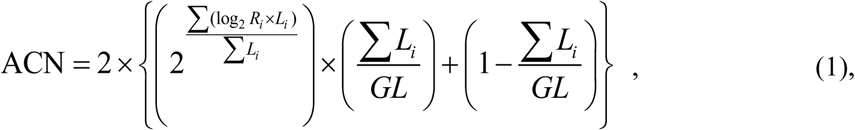

where log_2_*R*_*i*_, *L*_*i*_, and *GL* represent the log-ratio of CNA segment *i*, length of CNA segment *i*, and genome length (50 Gb for mouse, 40 Gb for human), respectively. CNAs of mouse bulk data were detected as previously described [41]. Briefly, segments were called for the same fore- and background BAM files as those used in SNV with Exome CNV [45] and Varscan2 [46]. Overlapping segments called by both tools were used as CNA segments.

### Random Forest

Random Forest was used to generate the classifier for the histological type and MSI status of human cancer. We used gene expression levels, number of SNVs in each gene, total mutation (SNV/indel) number, and mutation density (total number of SNVs/indels divided by target region size) as explanatory variables. Using TCGA data [24], we first filtered out unimportant explanatory variables using the two-sided Kruskal-Wallis test with *P* values of 5.00 × 10^-5^ and 1.00 × 10^−9^ yielding 171 and 78 variables for histological type and MSI status, respectively. These were used to train a Random Forest classifier with the “randomForest” function in the “randomForest” package of R software, with the options ntree = 10000 (setting the number of trees to grow to 1000) and mtry = 5 (setting the number of variables randomly sampled to five). Using the created classifier, the same explanatory variables for mouse data were used to classify each feature in the mouse model.

### MDS of mouse cell and TCGA samples

We first identified TCGA samples with gene expression patterns similar to the mouse single-cell groups. For that purpose, we calculated a normalized 1-*r* distance as follows:

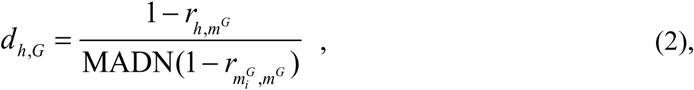

where *r*_*i,j*_ is a Pearson correlation coefficient between vectors *i* and *j* of expression levels in log across highly variable genes, *h* represents a human TCGA sample, *G* represents a mouse single-cell group, 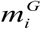 represents mouse single cell *i* in group *G, m*^*G*^ represents the centroid of 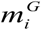 that was calculated by the median, and MADN represents the median absolute deviation adjusted by a factor for asymptotically normal consistency. We calculated this distance from a TCGA sample to every mouse group and selected a TCGA sample for those whose minimum distance across the groups was less than 4.05 and the difference between the first and second minimum distances was larger than 0.31. For selected TCGA and all mouse single-cell samples, MDS was performed based on the distance of 1-*r*.

## List of abbreviations

ITH: intra-tumor heterogeneity;
PCA: principal component analysis;
VAFs: variant allele frequencies;
MDS: multi-dimensional scaling;
SNV: single nucleotide variation;
TPM: transcripts per kilobase million;
LOWESS: locally weighted scatterplot smoothing;
ACN: average copy number.

## Declarations

### Ethics approval and consent to practice

Animal studies were carried out according to the Guideline for Animal Experiments established by the Committee for Ethics in Animal Experimentation of the National Cancer Center (T10-033-M05), which meets the ethical standards required by law and guidelines for animal experimentation in Japan.

### Consent for publication

Not applicable.

### Availability of data and materials

Sequence data used in this study are available in the DDBJ Sequenced Read Archive under Accession Nos. DRX100507-DRX100729. [These data are held until the acceptance. URL will be added after the data release.]

### Authors’ contributions

Y.A., Y.H., T.S., and M.K. designed the study. Y.A., T.M., T.I., and Y.H. performed the experiments. H.O., E.F., D.N., H.N., M.N., and M.K. analyzed the data. H.O., Y.A., E.F., T.I., Y.H., and M.K. wrote the manuscript. D. S., K.M., K.O., and T.S. reviewed the manuscript. M.K. led the project.

## Acknowledgements

We thank Joe Miyamoto, Asmaa Elzawahry, Iku Orihashi, Masako Ochiai, and Wakako Mukai for technical assistance, and Ryuichi Sugino and Daniel A. Vasco for useful suggestions.

