## Supplemental File 1 for "Single-cell DNA and RNA sequencing reveals the dynamics of intra-tumor heterogeneity in a colorectal cancer model"

### Supplementary Results

#### Estimation of false positive rate

We first estimated the number of SNV sites that differed between two individual mice of the pure C57BL/6J strain. For normal intestinal tract samples obtained from the two mice, we called SNVs in bulk-cell sequencing data using each of the two samples as the foreground data and the other as the background: the numbers were 1.0 and $4.5 \times 10^{-7}$ per chromosomal position for the two sample pairs, respectively. When we called SNVs in half-split sequencing data used as the fore- and background data for the same sample, the number of SNVs per position was 0 and $0.4 \times 10^{-7}$ for the two samples, respectively. Taken together, the false positive rate in bulk-cell sequencing was estimated as 1.0–4.9 ([1.0 + 0.0]–[4.5 + 0.4]) $\times 10^{-7}$. Because we called SNVs in single cells only at SNV sites called in bulk-cell sequencing data, the false-positive rate in single cells was not more than that in bulk-cell sequencing. Since 10–23% of chromosomal positions were called by our loose criteria for sequencing data from four single cells obtained from normal intestinal tract tissue, the false-positive rate per position in single-cell sequencing was estimated as 0.1–1.1 $\times 10^{-7}$.

#### Association with human cancers

We investigated the features of human colorectal cancer that correspond to those of our mouse cancer model using TCGA human colorectal cancer data and our mouse bulk sequencing data (39). We first examined individual molecular features. SNV density in the mouse model was closer to the hypermutation type of human colorectal cancer (Additional file 1: **Figure S7**). The expression of *MLH1*, the dysregulation of which causes hypermutation, was repressed with the levels decreasing over time (from T1 to T3) (Additional file 1: **Figure S7**). The average copy number across the mouse genome was closer to the hypermutation type, indicating low chromosomal instability (Additional file 1: **Figure S7**). Taken together, these results suggest that the mouse model was closer to the hypermutation type (albeit not extremely hyper) of human cancer.

We then analyzed clinical features in a machine learning approach (Random Forest) using a clinical feature as the objective variable and omics (SNV/indel/RNA) data as explanatory variables. Of the three histological types, including colon and rectal mucinous adenocarcinoma, our mouse model was closest to human colon adenocarcinoma and was closer to the MSI-high than MSI-low and microsatellite-stable types (Additional file 1: **Figure S8**). Thus, our mouse model represented the MSI-high hypermutation (although, not extremely hyper) type of human colon adenocarcinoma.

**Additional file 1: Supplementary Figures**

**A**

| HumanGene | Chr | Start | End | Mut_type | Ref | Alt | Notion | Reference |
|---|---|---|---|---|---|---|---|---|
| *KRAS* | 6 | 145,169,253 | 145,169,253 | indel | - | A | intronic | TCGA SMG |
| *TP53* | 11 | 69,402,151 | 69,402,151 | snv | A | T | nonsynonymous SNV | TCGA SMG and COSMIC |
| *CLTC* | 11 | 86,520,656 | 86,520,656 | snv | A | T | nonsynonymous SNV | COSMIC |
| *ALK* | 17 | 72,952,883 | 72,952,883 | snv | A | C | nonsynonymous SNV | COSMIC |
| *LRP1B* | 2 | 40,724,718 | 40,724,718 | snv | C | T | nonsynonymous SNV | COSMIC |
| *GRIN2A* | 16 | 9,579,188 | 9,579,188 | snv | T | C | nonsynonymous SNV | COSMIC |
| *MSH2* | 17 | 88,079,144 | 88,079,144 | snv | C | T | nonsynonymous SNV | COSMIC |
| *SALL4* | 2 | 168,580,005 | 168,580,005 | snv | C | T | nonsynonymous SNV | COSMIC |

**B**

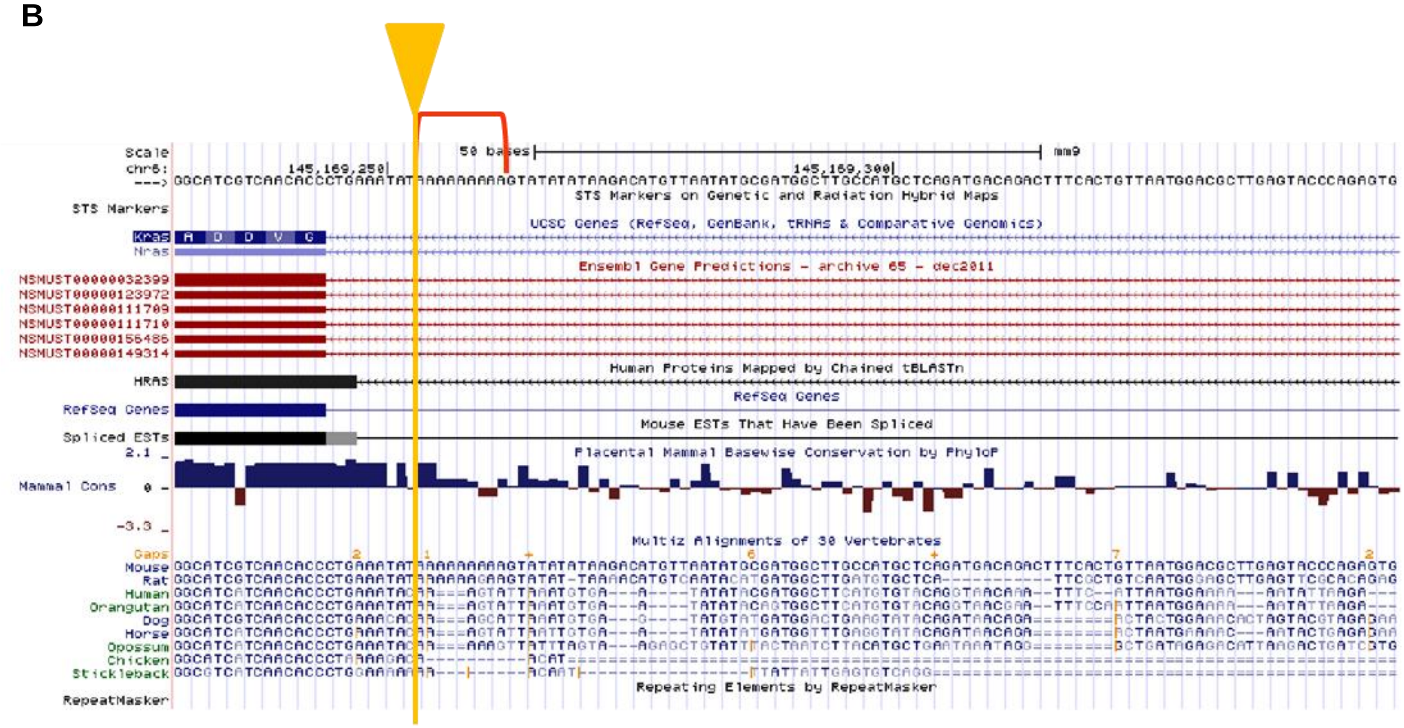


**Figure S1** Details of mutations found in the significantly mutated genes of TCGA colorectal cancer and in cancer-related genes referred in COSMIC by bulk-cell DNA sequencing.

(**A**) Annotations of genes found in the significantly mutated genes of TCGA colorectal cancer and in COSMIC cancer-related genes by bulk-cell DNA sequencing. (**B**) The *KRAS* mutation in the mouse genome by the UCSC genome browser. The reversed U symbol in red indicates a mono-repeat of A. The arrow and line in gold indicate the position of mutation.

**A**

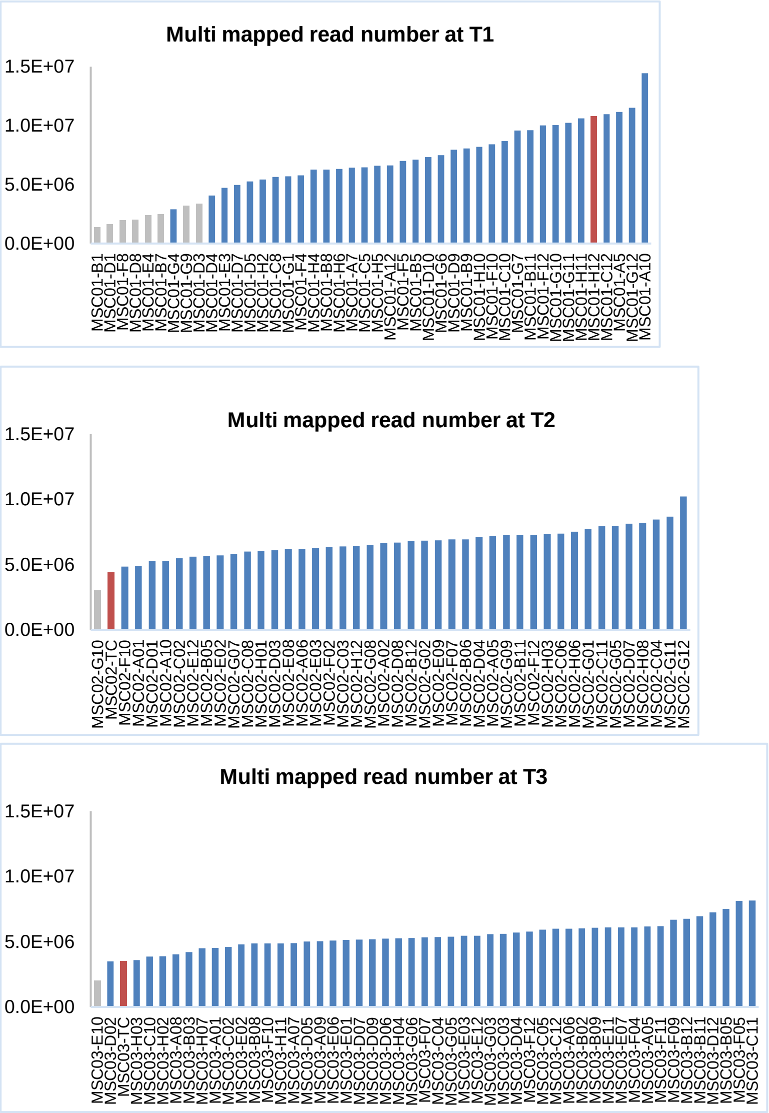

Multi mapped read number at T1
1.5E+07
1.0E+07
5.0E+06
0.0E+00
Multi mapped read number at T2
1.5E+07
1.0E+07
5.0E+06
0.0E+00
Multi mapped read number at T3
1.5E+07
1.0E+07
5.0E+06
0.0E+00

**B**

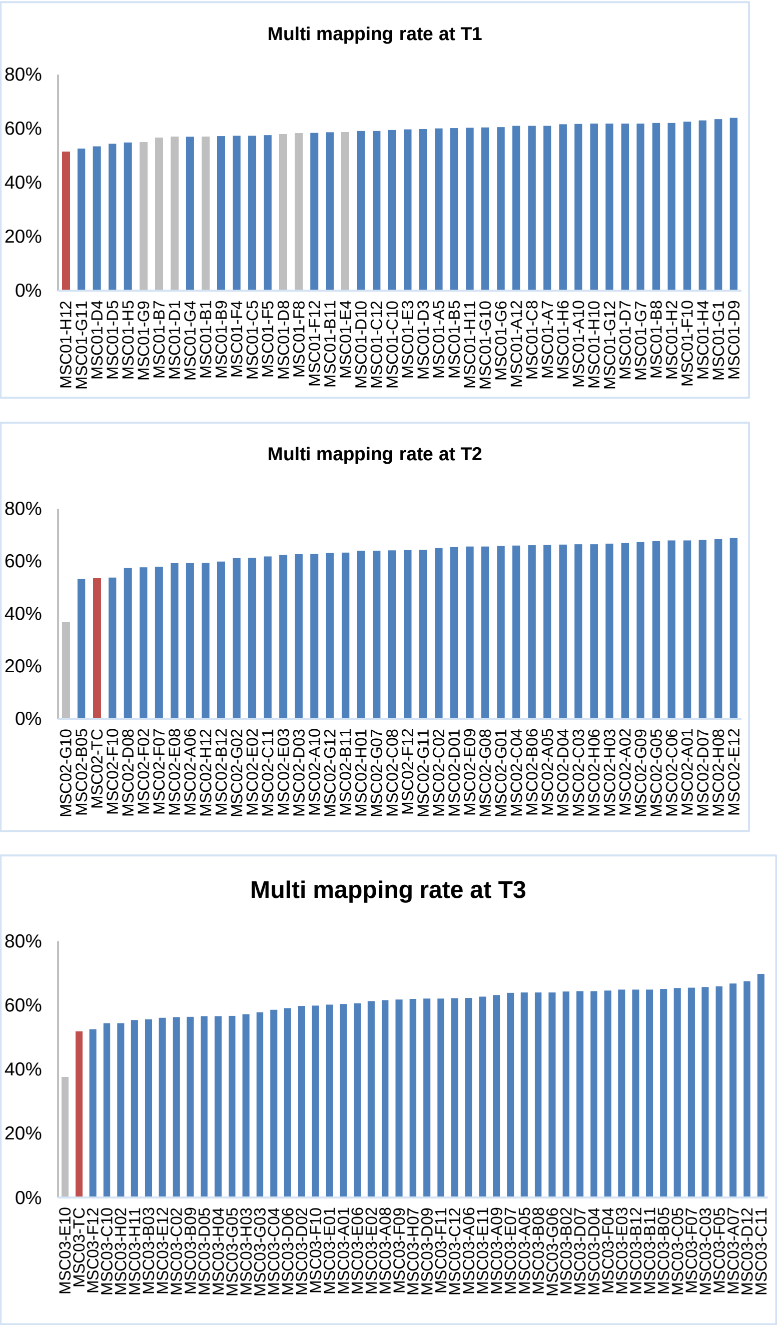

Multi mapping rate at T1
80%
60%
40%
20%
0%
MSC01-H12
MSC01-G11
MSC01-D4
MSC01-D5
MSC01-H5
MSC01-G9
MSC01-B7
MSC01-D1
MSC01-G4
MSC01-B1
MSC01-B9
MSC01-F4
MSC01-C5
MSC01-F5
MSC01-D8
MSC01-F8
MSC01-F12
MSC01-B11
MSC01-E4
MSC01-D10
MSC01-C12
MSC01-C10
MSC01-E3
MSC01-D3
MSC01-A5
MSC01-B5
MSC01-H11
MSC01-G10
MSC01-G6
MSC01-A12
MSC01-C8
MSC01-A7
MSC01-H6
MSC01-A10
MSC01-H10
MSC01-G12
MSC01-D7
MSC01-G7
MSC01-B8
MSC01-H2
MSC01-F10
MSC01-H4
MSC01-G1
MSC01-D9
Multi mapping rate at T2
80%
60%
40%
20%
0%
MSC02-G10
MSC02-B05
MSC02-TC
MSC02-F10
MSC02-D08
MSC02-F02
MSC02-F07
MSC02-E08
MSC02-A06
MSC02-H12
MSC02-B12
MSC02-G02
MSC02-E02
MSC02-C11
MSC02-F05
MSC02-E03
MSC02-D03
MSC02-A10
MSC02-G12
MSC02-B11
MSC02-H01
MSC02-G07
MSC02-C08
MSC02-F12
MSC02-G11
MSC02-C02
MSC02-D01
MSC02-E09
MSC02-G08
MSC02-G01
MSC02-C04
MSC02-B06
MSC02-A05
MSC02-D04
MSC02-C03
MSC02-H06
MSC02-H03
MSC02-A02
MSC02-G09
MSC02-G05
MSC02-C06
MSC02-A01
MSC02-D07
MSC02-H08
MSC02-E12
Multi mapping rate at T3
80%
60%
40%
20%
0%
MSC03-E10
MSC03-TC
MSC03-F12
MSC03-C10
MSC03-H02
MSC03-H11
MSC03-B03
MSC03-E12
MSC03-C02
MSC03-B09
MSC03-D05
MSC03-H04
MSC03-G05
MSC03-H03
MSC03-G03
MSC03-C04
MSC03-D06
MSC03-D02
MSC03-F10
MSC03-E01
MSC03-A01
MSC03-E06
MSC03-E02
MSC03-A08
MSC03-F09
MSC03-H07
MSC03-D09
MSC03-F11
MSC03-C12
MSC03-A06
MSC03-E11
MSC03-A09
MSC03-E07
MSC03-A05
MSC03-B08
MSC03-G06
MSC03-B02
MSC03-D07
MSC03-D04
MSC03-F04
MSC03-E03
MSC03-B12
MSC03-B11
MSC03-B05
MSC03-C05
MSC03-F07
MSC03-C03
MSC03-F05
MSC03-A07
MSC03-D12
MSC03-C11

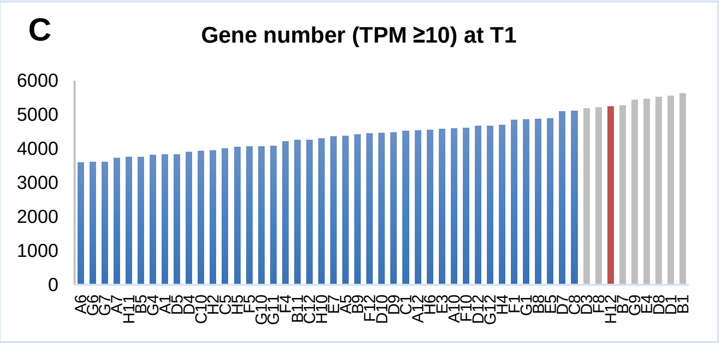
C
Gene number (TPM ≥10) at T1
6000
5000
4000
3000
2000
1000
0

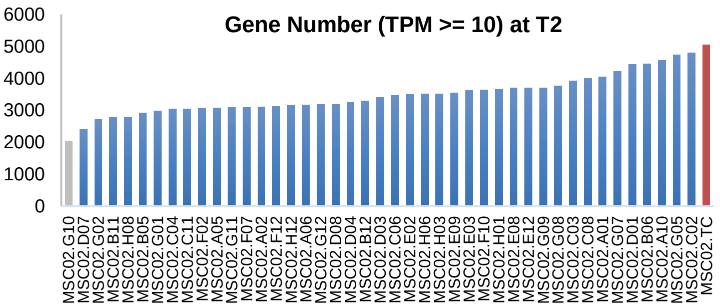
Gene Number (TPM >= 10) at T2
6000
5000
4000
3000
2000
1000
0

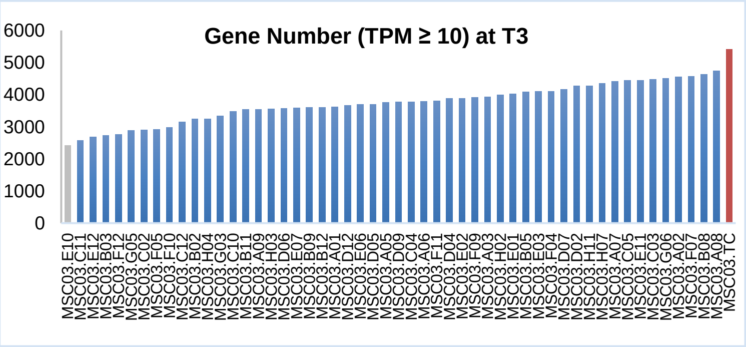
Gene Number (TPM ≥ 10) at T3
6000
5000
4000
3000
2000
1000
0

D

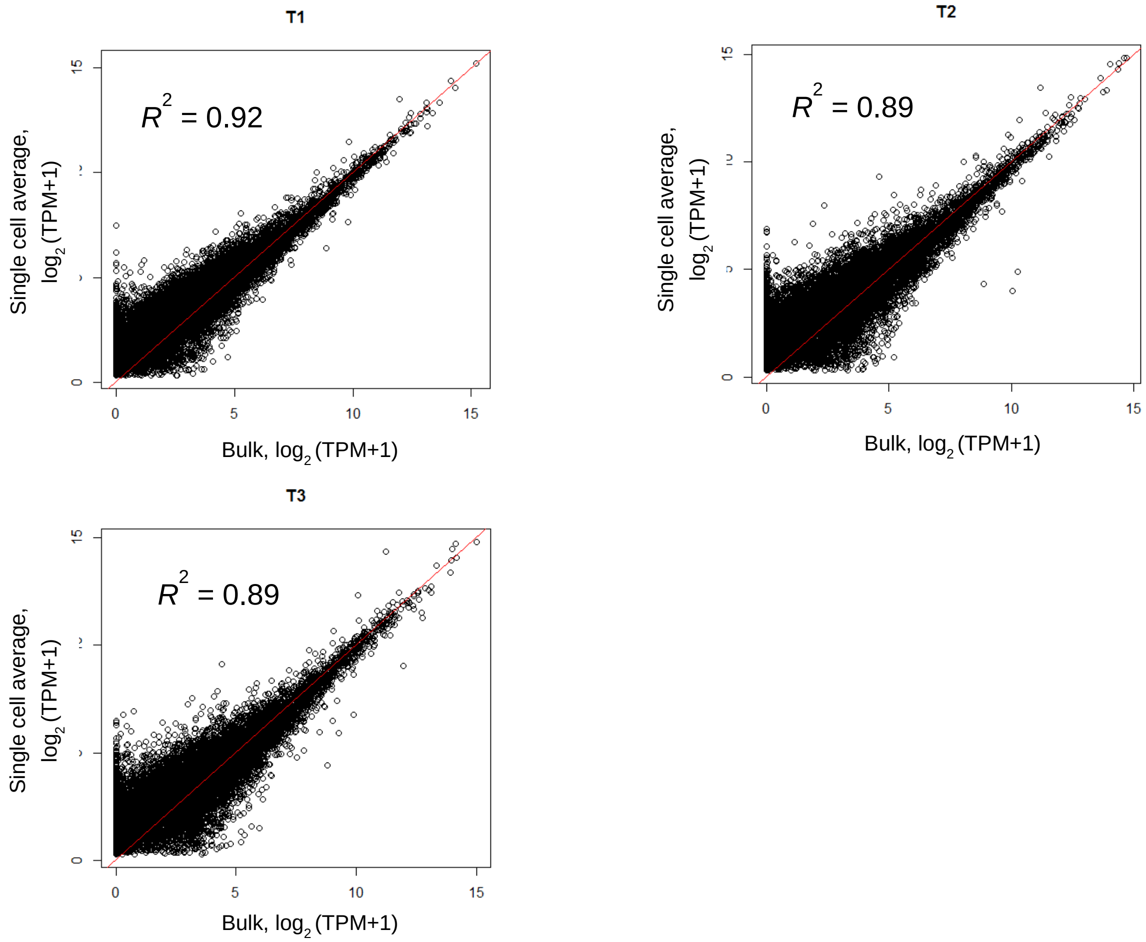

T1
$R^2$ = 0.92
Single cell average, log2 (TPM+1)
Bulk, log2 (TPM+1)
0
5
10
15
T2
$R^2$ = 0.89
Single cell average, log2 (TPM+1)
Bulk, log2 (TPM+1)
T3
$R^2$ = 0.89
Single cell average, log2 (TPM+1)
Bulk, log2 (TPM+1)


E

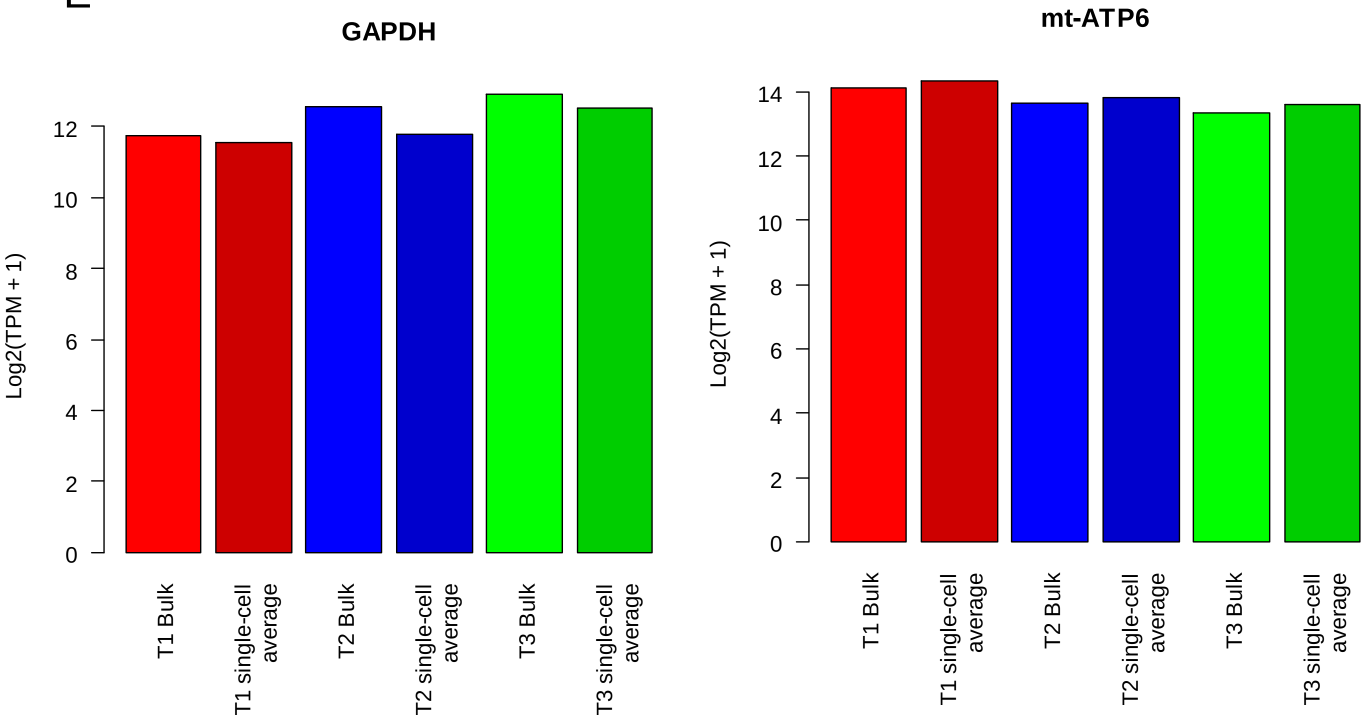

GAPDH
Log2(TPM + 1)
0
2
4
6
8
10
12
T1 Bulk
T1 single-cell average
T2 Bulk
T2 single-cell average
T3 Bulk
T3 single-cell average
mt-ATP6
Log2(TPM + 1)
14

**Figure S2** Quality check of single-cell transcriptome sequencing data. (A) Number of mapped reads, (B) mapping rate, and (C) number of expressed genes (TPM $\geq$ 10). We removed outliers (gray) based on the combination of the number of expressed genes ($\leq$5200) and number of mapped genes ($\leq 2.2 \times 10^6$), and mapping rate ($\leq$20%). Blue and orange bars represent single-cell samples that were ultimately used and bulk samples, respectively. (D) Scatter plot of gene expression levels from a bulk sample versus expression levels averaged across the single cells that were ultimately used. (E) The expression levels of housekeeping genes across T1, T2, and T3.

A

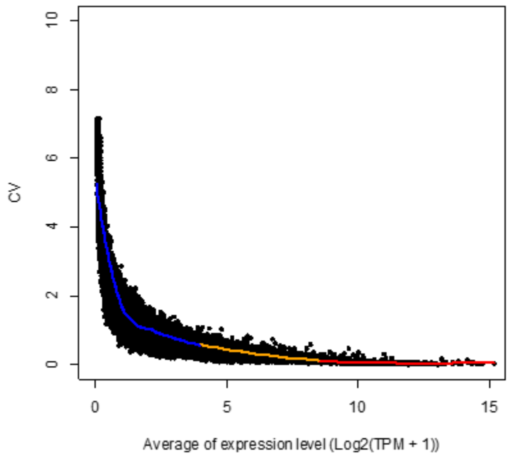
CV
Average of expression level (Log2(TPM + 1))

B

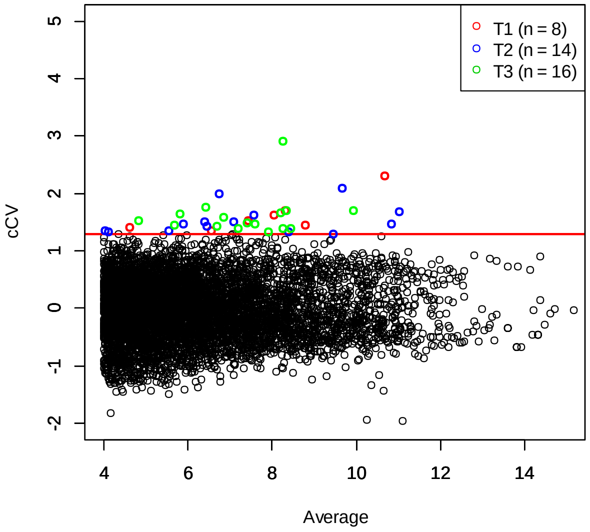
T1 (n = 8)
T2 (n = 14)
T3 (n = 16)
cCV
Average

C

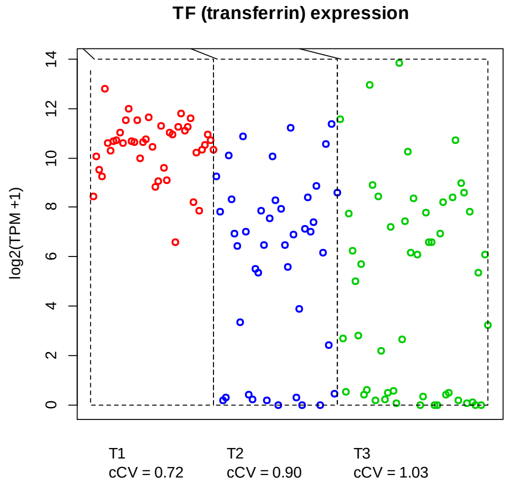
TF (transferrin) expression
log2(TPM +1)
T1
cCV = 0.72
T2
cCV = 0.90
T3
cCV = 1.03

**Figure S3** c*CV* and highly variable genes. (A) *CV* versus gene expression levels averaged across single cells. Regression analysis was performed to obtain the locally weighted scatterplot smoothing (LOWESS) curve within the range indicated by each color (blue, yellow, and red). (B) c*CV* and average expression levels. Highly variable genes are shown above the red line. (C) c*CV* and distribution of gene expression levels across single cells, illustrated with the transferrin gene. Each circle represents gene expression level in a single cell.

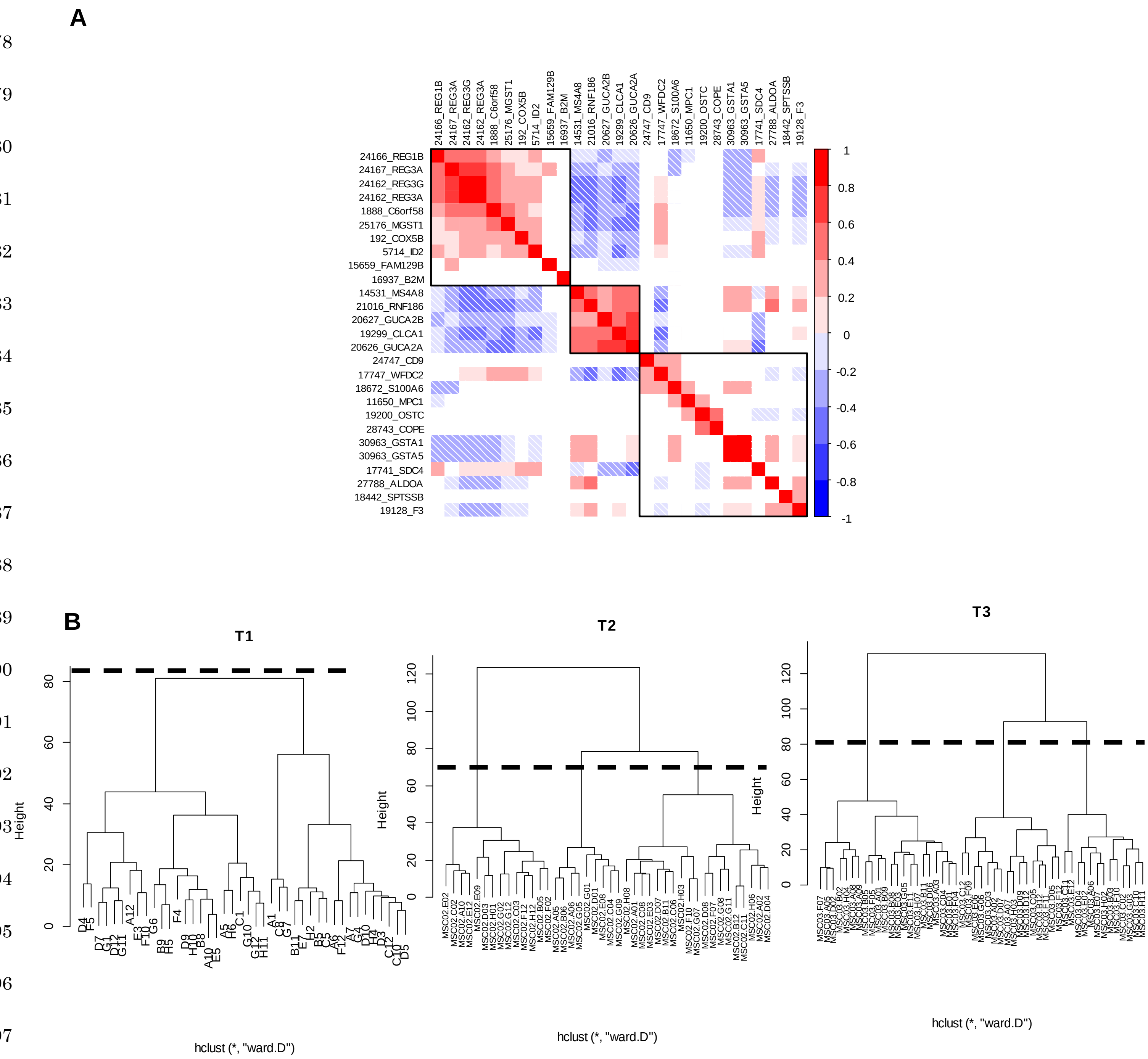


**Figure S4** Determination of gene and cell groups in single-cell RNA sequencing. (A) Correlation plot of highly variable genes. (B) Dendrogram of single cells.

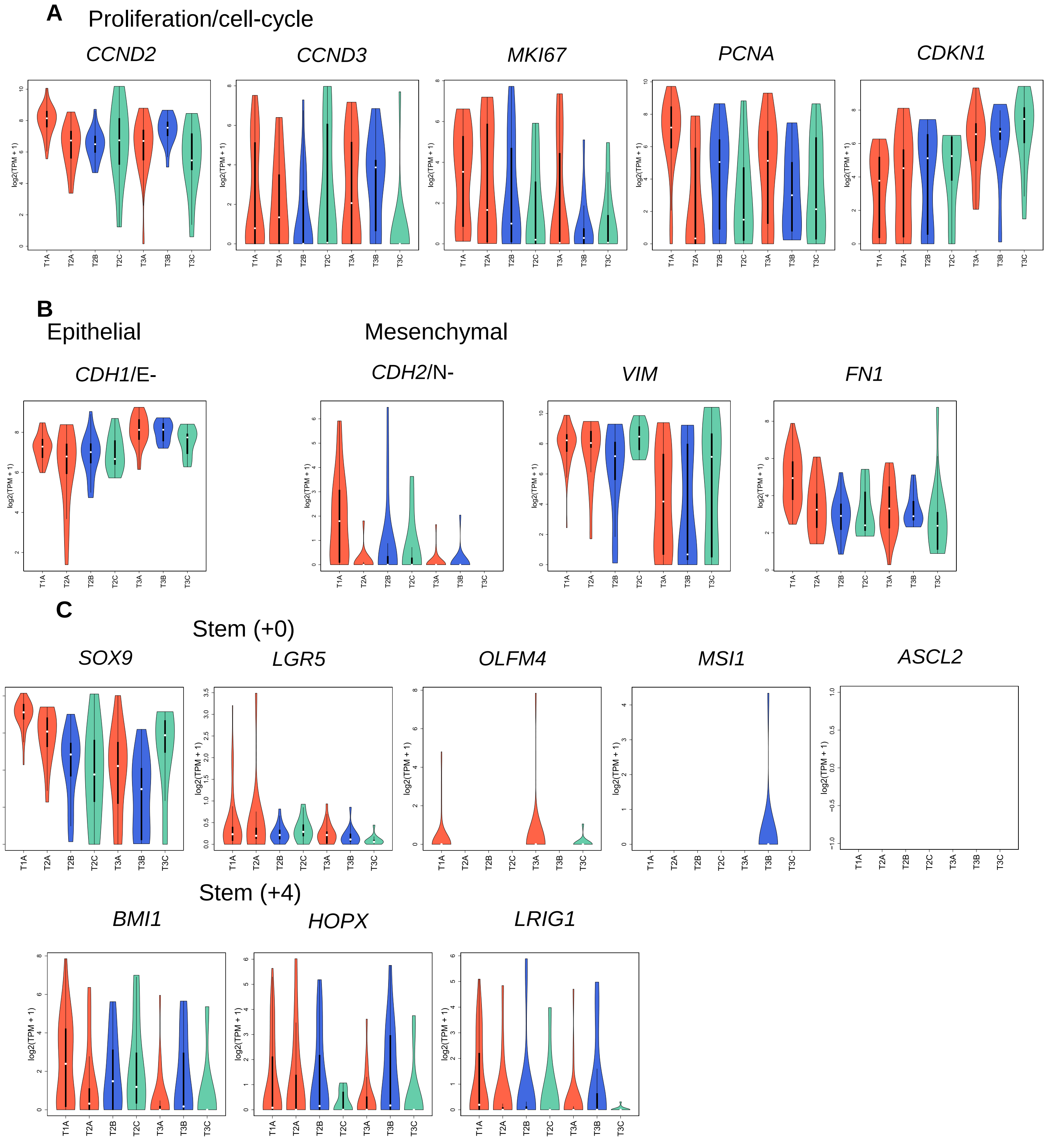
A
Proliferation/cell-cycle
CCND2
CCND3
MKI67
PCNA
CDKN1
B
Epithelial
Mesenchymal
CDH1/E-
CDH2/N-
VIM
FN1
C
Stem (+0)
SOX9
LGR5
OLFM4
MSI1
ASCL2
Stem (+4)
BMI1
HOPX
LRIG1
log2(TPM + 1)
T1A
T2A
T2B
T2C
T3A
T3B
T3C

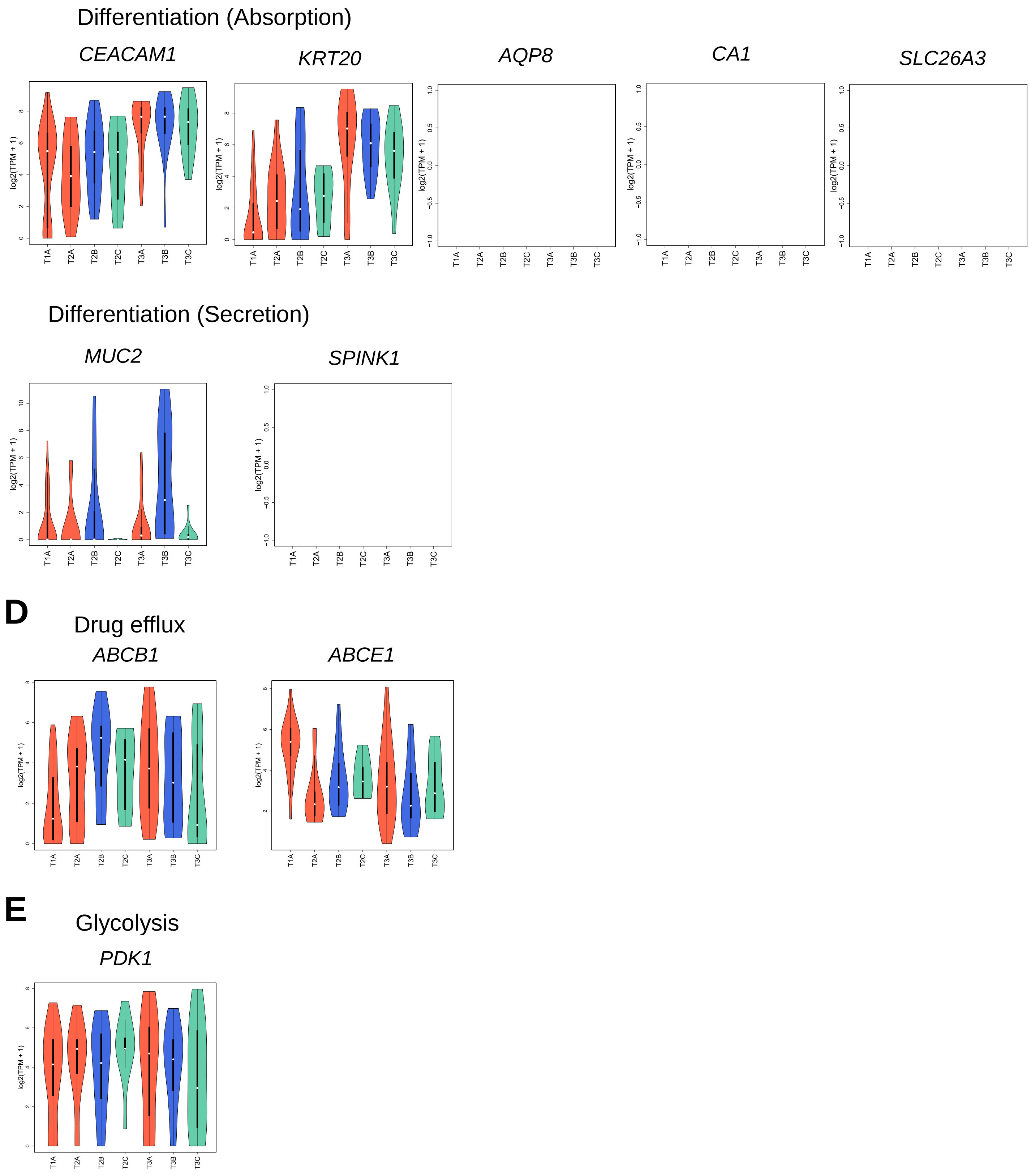
Differentiation (Absorption)
CEACAM1
KRT20
AQP8
CA1
SLC26A3
log2(TPM + 1)
T1A
T2A
T2B
T2C
T3A
T3B
T3C
Differentiation (Secretion)
MUC2
SPINK1
D
Drug efflux
ABCB1
ABCE1
E
Glycolysis
PDK1

**Figure S5** Violin plots of the expression levels of the marker genes. “A,” “B,” and “C” followed by T1/T2/T3 represent Anti-Epithelial, cGMP/GC, and Dormant cell groups, respectively (*n*: 42 for T1A, 14 for T2A, 19 for T2B, 9 for T2C, 22 for T3A, 16 for T3B, and 13 for T3C). Some genes such as *ASCL2* were not expressed in any category. (A) Proliferation/cell-cycle markers, (B) epithelial and mesenchymal markers, (C) stem cell and differentiation markers, (D) drug efflux markers, and (E) glycolysis markers.

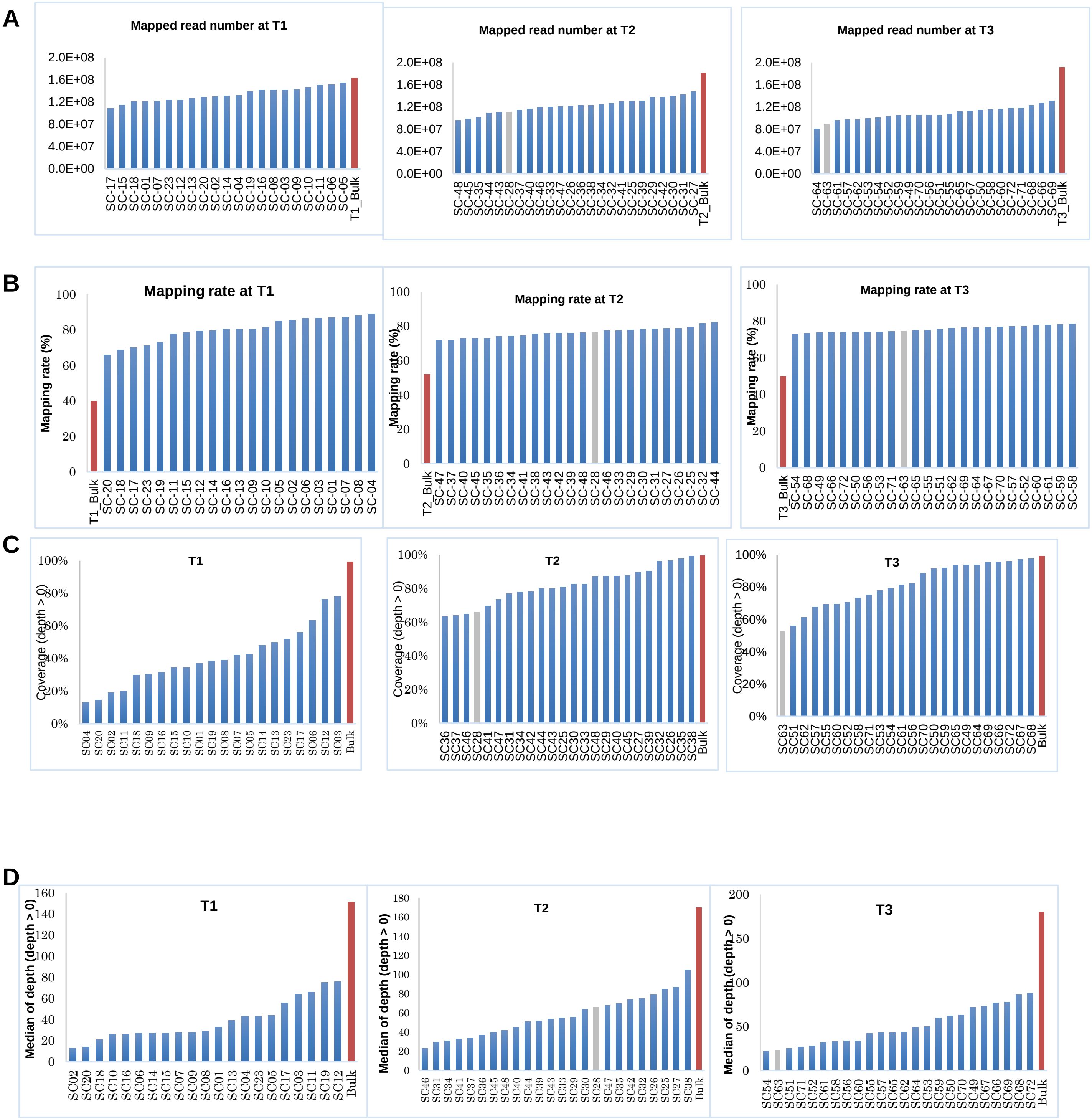
A
Mapped read number at T1
Mapped read number at T2
Mapped read number at T3
B
Mapping rate at T1
Mapping rate at T2
Mapping rate at T3
Mapping rate (%)
C
T1
T2
T3
Coverage (depth > 0)
D
T1
T2
T3
Median of depth (depth > 0)

E

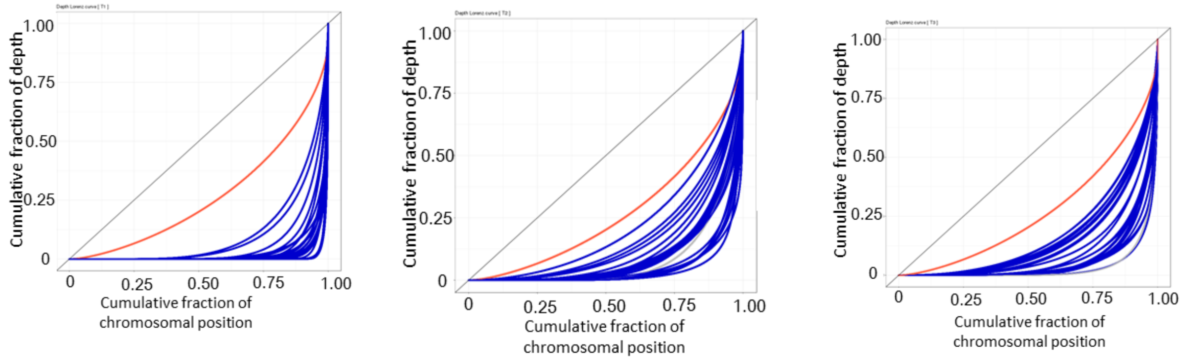
Cumulative fraction of depth
Cumulative fraction of chromosomal position
0
0.25
0.50
0.75
1.00

F

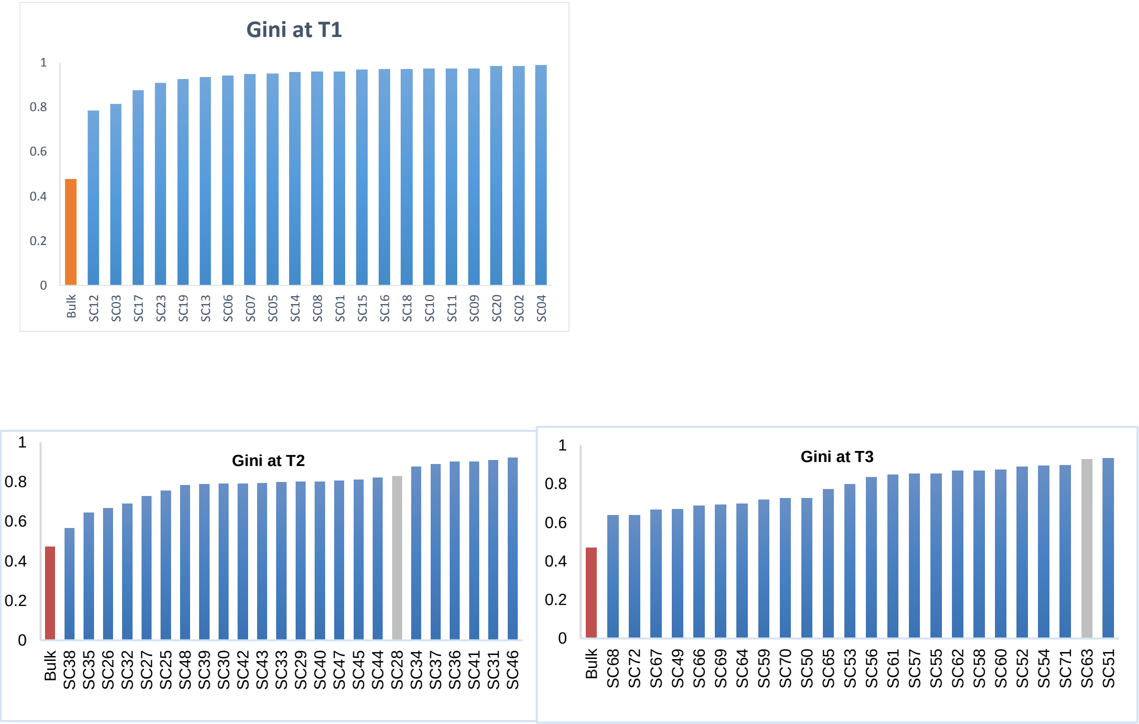
Gini at T1
Gini at T2
Gini at T3
Bulk
SC12
SC03
SC17
SC23
SC19
SC13
SC06
SC07
SC05
SC14
SC08
SC01
SC15
SC16
SC18
SC10
SC11
SC09
SC20
SC02
SC04
SC38
SC35
SC26
SC32
SC27
SC25
SC48
SC39
SC30
SC42
SC43
SC33
SC29
SC40
SC47
SC45
SC44
SC28
SC34
SC37
SC36
SC41
SC31
SC46
SC68
SC72
SC67
SC49
SC66
SC69
SC64
SC59
SC70
SC50
SC65
SC53
SC56
SC61
SC57
SC55
SC62
SC58
SC60
SC52
SC54
SC71
SC63
SC51

**G**

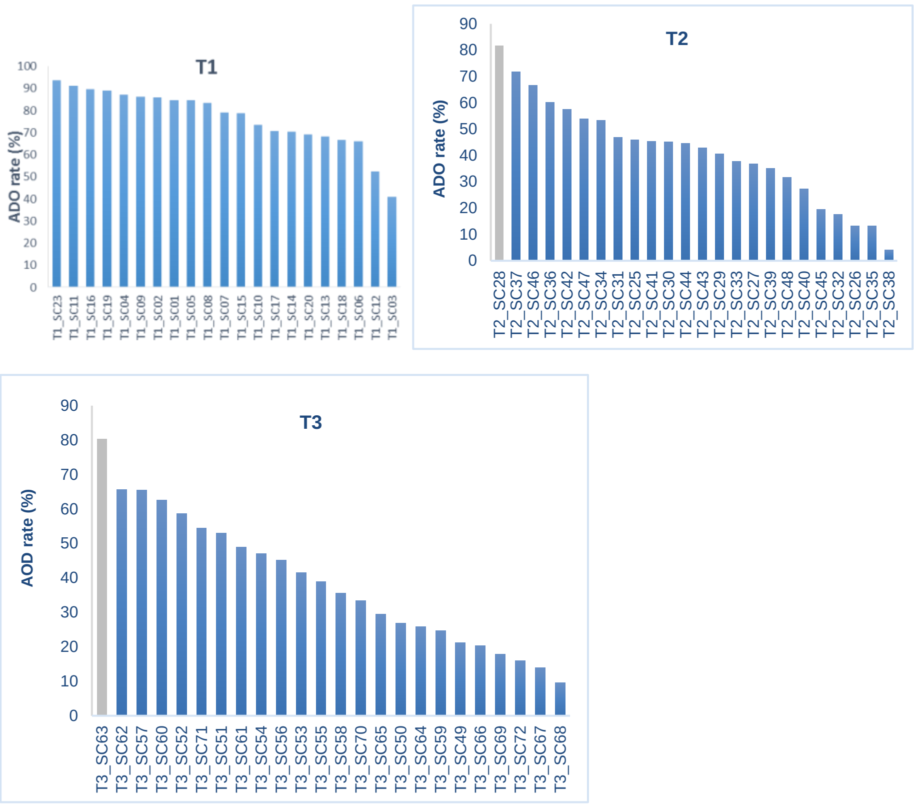


**H**

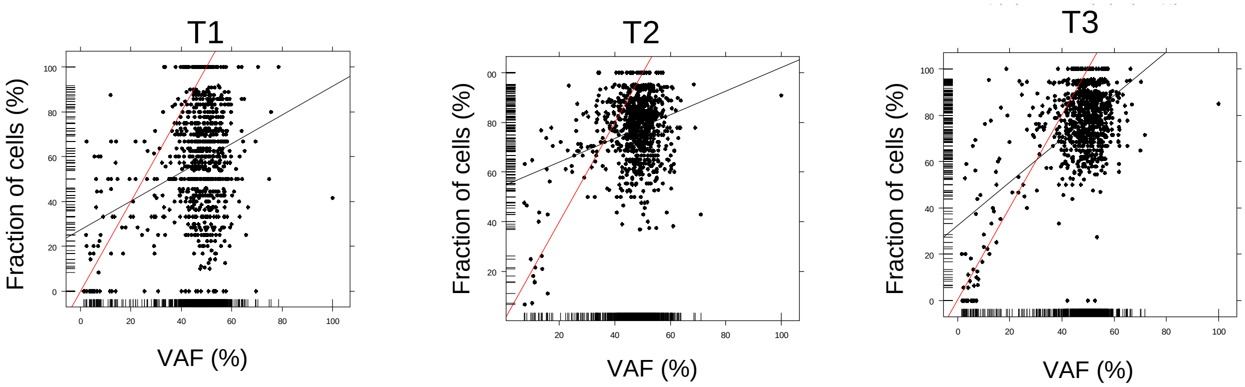


**Figure S6** Quality check of single-cell exome sequencing data. Bars/curves in orange, blue, and gray represent bulk-cell, single-cell, and filtered-out data, respectively. Shown are the (A) number of mapped reads; (B) mapping rate; (C) coverage of genome with depth > 0; (D) median depth, in which regions with depth = 0 were excluded; (E) Lorenz curve of depth (including regions with depth = 0); (F) Gini coefficients of depth (including regions with depth = 0); (G) ADO rate; and (H) Scatter plot of SNVs between VAFs in bulk-cell sequencing and fractions of single cells with SNVs called in single-cell sequencing. Black and red lines represent the linear regression and theoretically expected lines, respectively.

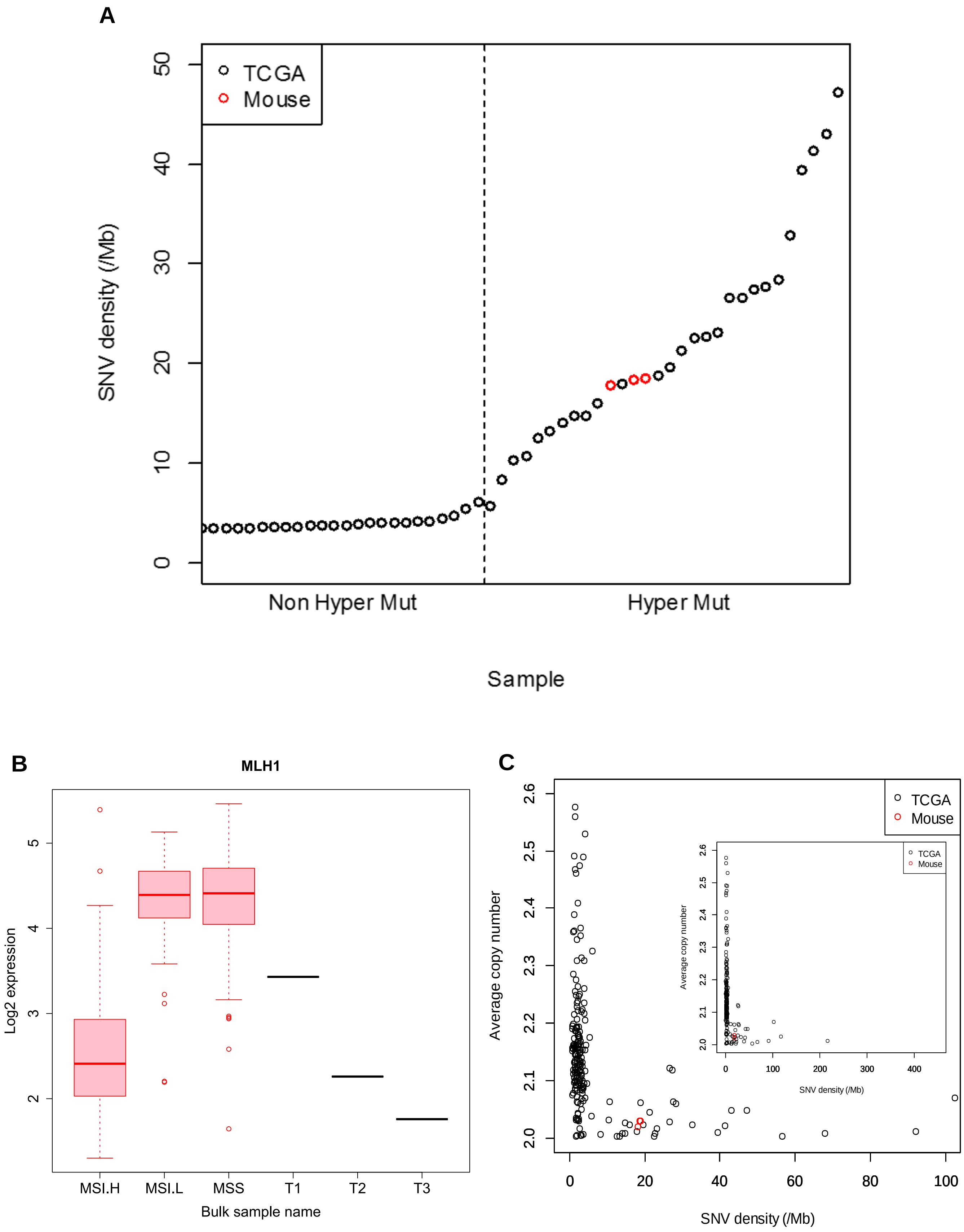
A
TCGA
Mouse
SNV density (/Mb)
0
10
20
30
40
50
Non Hyper Mut
Hyper Mut
Sample
B
MLH1
Log2 expression
2
3
4
5
MSI.H
MSI.L
MSS
T1
T2
T3
Bulk sample name
C
TCGA
Mouse
Average copy number
2.0
2.1
2.2
2.3
2.4
2.5
2.6
0
20
40
60
80
100
SNV density (/Mb)
0
100
200
300
400

**Figure S7** Human cancer counterpart to our mouse model according to molecular features. (A) SNV density in human colorectal cancer and in the mouse model. Black and red circles represent TCGA human colorectal cancer samples and mouse samples at T1, T2, and T3, respectively. Dashed lines separate hyper and non-hyper mutation types. (B) *MLH1* expression in TCGA and mouse samples. MSI.H, microsatellite instability high ($n$ = 35); MSI.L, microsatellite instability low ($n$ = 42); MSS, microsatellite stable ($n$ = 166). (C) Average copy number across the genome versus SNV density. Insets in panels A and C show zoomed-out views.

A

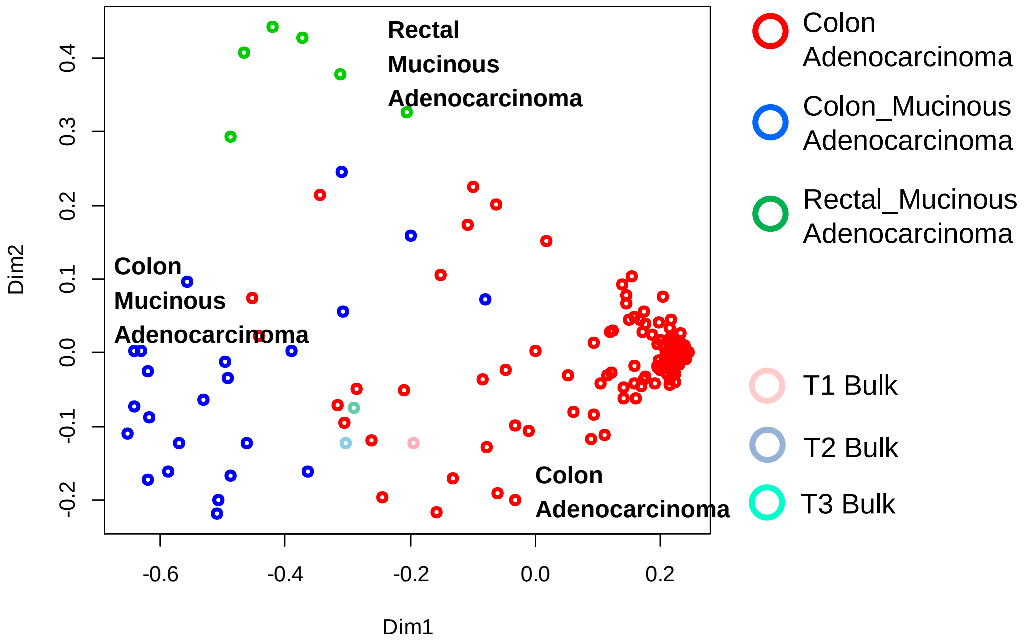


B

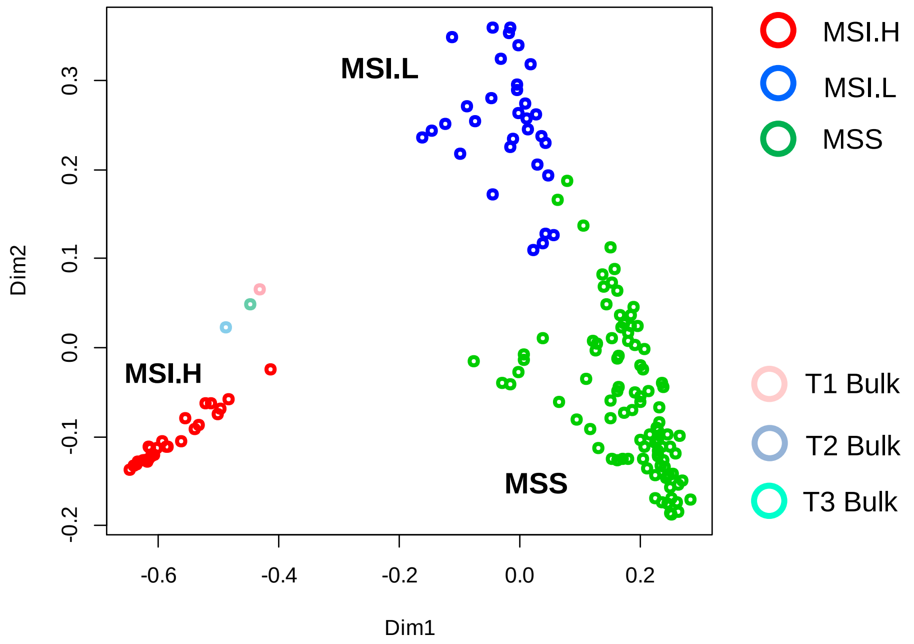


**Figure S8** Human cancer counterpart to our mouse model according to clinical features. Multidimensional scaling plots generated by Random Forest based on the proximity matrix are shown. (A) For histological type. (B) For microsatellite instability. MSI.H, microsatellite instability high; MSI.L, microsatellite instability low; MSS, microsatellite stable.

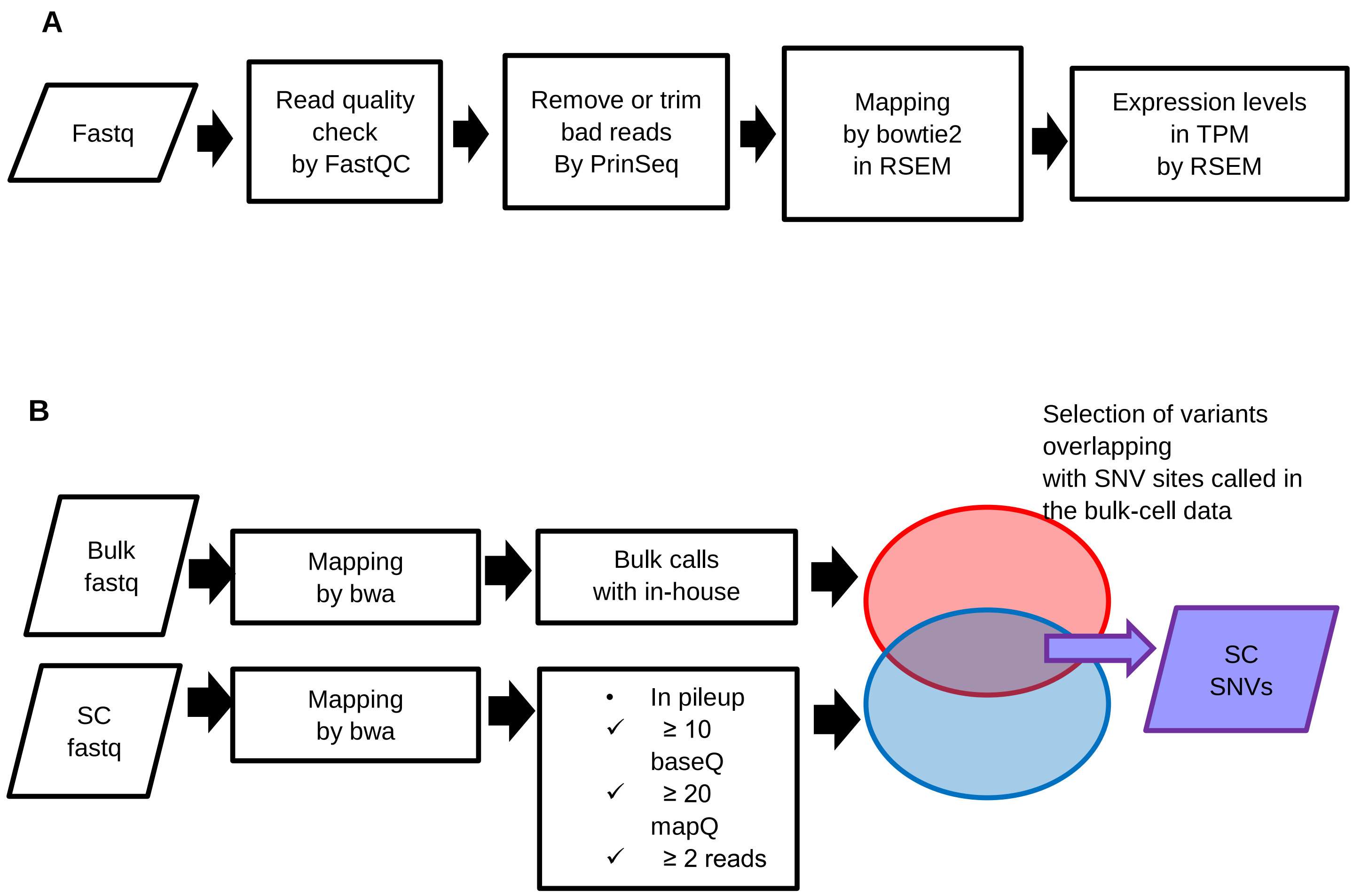


**Figure S9** Procedure for calculating expression levels and for calling SNVs in single-cell sequencing. (A) Procedure for calculating expression levels (TPM). (B) Procedure for calling SNVs in single cells (SCs).
